# EZH2 synergizes with BRD4-NUT to drive NUT carcinoma growth through silencing of key tumor suppressor genes

**DOI:** 10.1101/2023.08.15.553204

**Authors:** Yeying Huang, R. Taylor Durall, Nhi M. Luong, Hans J. Hertzler, Julianna Huang, Prafulla C. Gokhale, Brittaney A. Leeper, Nicole S. Persky, David E. Root, Praju V. Anekal, Paula D.L.M. Montero Llopis, Clement N. David, Jeffery L. Kutok, Alejandra Raimondi, Karan Saluja, Jia Luo, Cynthia A. Zahnow, Biniam Adane, Kimberly Stegmaier, Catherine E. Hawkins, Christopher Ponne, Quan Le, Geoffrey I. Shapiro, Madeleine E. Lemieux, Kyle P. Eagen, Christopher A. French

## Abstract

NUT carcinoma (NC) is an aggressive carcinoma driven by the BRD4-NUT fusion oncoprotein, which activates chromatin to promote expression of pro-growth genes. BET bromodomain inhibitors (BETi) impede BRD4-NUT’s ability to activate genes and are thus a promising treatment but limited as monotherapy. The role of gene repression in NC is unknown. Here, we demonstrate that EZH2, which silences genes through establishment of repressive chromatin, is a dependency in NC. Inhibition of EZH2 with the clinical compound tazemetostat (taz) potently blocked growth of NC cells. Epigenetic and transcriptomic analysis revealed that taz reversed the EZH2-specific H3K27me3 silencing mark, and restored expression of multiple tumor suppressor genes while having no effect on key oncogenic BRD4- NUT-regulated genes. *CDKN2A* was identified as the only gene amongst all taz-derepressed genes to confer resistance to taz in a CRISPR-Cas9 screen. Combined EZH2 inhibition and BET inhibition synergized to downregulate cell proliferation genes resulting in more pronounced growth arrest and differentiation than either inhibitor alone. In pre-clinical models, combined taz and BETi synergistically blocked growth and prolonged survival of NC-xenografted mice, with all mice cured in one cohort.

**STATEMENT OF SIGNIFICANCE:** Identification of EZH2 as a dependency in NC substantiates the reliance of NC tumor cells on epigenetic dysregulation of functionally opposite, yet highly complementary chromatin regulatory pathways to maintain NC growth. In particular, repression of CDKN2A expression by EZH2 provides a mechanistic rationale for combining EZH2i with BETi for the clinical treatment of NC.

## INTRODUCTION

NUT carcinoma (NC) is an aggressive squamous carcinoma driven most commonly (78%) by the BRD4- NUT fusion oncoprotein^1^. NC is the second-most aggressive solid tumor in humans^2, 3^, with a median survival of 6.5 months^1^. It affects patients of all ages, but is most common in adolescents and young adults, often referred to as the AYA gap. The estimated annual U.S. incidence of NC, based on sequential next generation DNA sequencing (NGS)^4^, is roughly 0.06% of all solid tumors, or by extrapolation 1,100 cases. NC accounts for a substantial portion of poorly differentiated carcinomas^5, 6^, and its diagnosis is rapidly increasing with both awareness and ease of diagnostic testing^7^. Currently, there are no effective systemic treatments for NC, making it a disease with an extreme unmet need^6^.

Mechanistically, BRD4 is a member of the BET family of proteins, which bind acetyl-histones through their dual bromodomains. Upon fusion to NUT, which recruits and stimulates the histone acetyltransferase p300, BRD4-NUT forms enormous, H3K27ac-enriched, super-enhancers called ‘megadomains’. Megadomains maintain expression of key oncogenic genes, *MYC* and *SOX2*^8, 9^. BRD4- NUT also reprograms chromatin 3D structure resulting in inter-chromosomal DNA interactions between these lineage-defining transcription factors^10, 11^. Megadomains and chromatin 3D interactions mediated by BRD4-NUT can be visualized immunohistochemically by light microscopy or by immunofluorescence microscopy^8, 12–14^. The functional outcome of these BRD4-NUT-driven processes is blockade of differentiation and maintenance of growth of NC cells. When BRD4-NUT or variant NUT-fusion proteins are depleted, NC cells rapidly arrest growth and undergo terminal squamous differentiation and stop proliferating^15, 16^.

BET bromodomain inhibitors (BETi), small molecule acetyl-histone mimetics, evict BRD4-NUT from chromatin, arrest NC growth, and induce differentiation^17^. As *BRD4-NUT* is the only known *BRD4* oncogene, on-target activity of BETi in NC patients^18^ led to a new field investigating the role of BRD4 in numerous malignancies and the application of BETi as a treatment option for many cancers^19^. However, it has become clear that BETi monotherapy does not fully address NC biology. In clinical trials, the best responses of NC to BETi are short-lived (1-3 months), partial responses in 20-30% of patients^6, 20, 21^.

The lack of other oncogenic driver mutations in NC^1, 18, 22, 23^, and its occurrence in infants, have led us to believe that BRD4-NUT (and variants) act alone to transform and maintain growth of NC. However, mechanisms that evade growth suppression, well-known in other cancers^24^, have not been identified for NC. That BRD4-NUT promotes gene expression prompted us to investigate epigenetic gene silencing as a parallel pathway that regulates NC growth. We find that BRD4-NUT-mediated gene activation is highly complemented by silencing of tumor suppressor genes. Based on phenotypic, transcriptomic, and epigenomic analysis, we identify a new vulnerability in NC that can be pharmacologically modulated. Identification of this obligate dependency enabled us to develop and *in vivo* validate a synergistic drug combination with strong clinical potential for the treatment of NC.

## RESULTS

### EZH2 is required for maintenance of growth and blockade of differentiation of NC

We reasoned that facultative heterochromatin may be repressing the expression of genes that suppress growth in NC. We therefore focused on Polycomb-group proteins that are canonical gene repressors. To examine Polycomb-group protein expression levels from clinical samples of NC we created a tumor microarray (TMA). Our TMA consists of 77 different NCs with four 1mm cores from each tumor. Two cores were taken from the tumor center and two from the tumor margin for a total of 278 cores. We performed immunohistochemistry (IHC) for Enhancer of Zeste Homolog 2 (EZH2), the catalytic subunit of polycomb repressive complex 2 (PRC2), which catalyzes histone H3K27me3. 84% of cases (58 of 69 stainable tumors) demonstrated high EZH2 expression, defined by ≥ 50% nuclear staining (**Fig. 1A**).

**Fig. 1.**
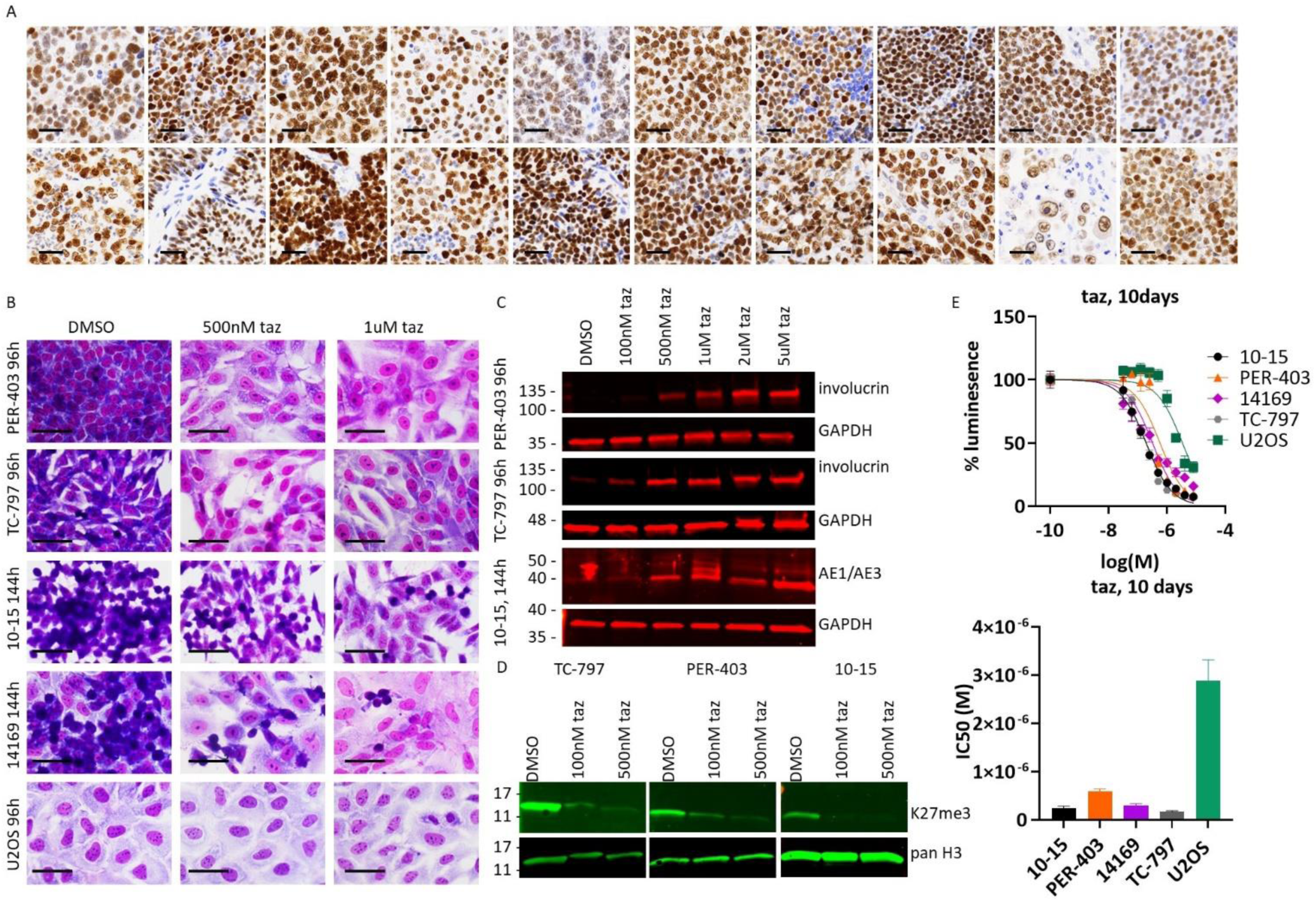
EZH2 is required for maintenance of growth and blockade of differentiation of NC. A. Immunohistochemistry (IHC) for EZH2 in 20 different NUT carcinoma primary tumors. Scale bar, 25µm. B. Hemacolor stained NC cell lines grown on coverslips. Scale bar, 25µm. C. Immunoblots of indicated proteins. Imaged on a Li-Cor Odyssey CLx fluorescence gel imager. D. Immunoblots of indicated histones and histone post-translational modifications. H3, histone H3; K27me3, histone H3 K27me3. E. Dose response curves from representative single replicates using Cell Titer Glo as readout. Below, IC50s from biological triplicate, technical quintuplet, dose-response assays corresponding with the above.

Aberrant EZH2 function has been implicated in many cancers^25^, and because NC is an epigenetic malignancy, we reasoned it might also be involved in NC pathogenesis. Although there is no evidence of oncogenic *EZH2* mutations in NC, based on reported whole or targeted genome sequencing of at least 21 primary tumors^1, 18, 26^ and 15 cell lines^22, 23^, robust EZH2 expression led us to evaluate the role of EZH2-mediated H3K27me3 in NC. A clinical compound, tazemetostat (taz, EPZ-6438), inhibits EZH2’s H3K27 methyltransferase activity. Taz has been FDA-approved for treatment of epithelioid sarcoma and has accelerated FDA approval in follicular lymphoma^27^. We thus evaluated the effects of EZH2 catalytic inhibition by taz on NC growth and differentiation. Reversal of H3K27me3 by taz is slow, occurring over days, due to the slow kinetics of H3K27 demethylation^28, 29^, and the phenotypic effects are not fully realized until 10-14 days ^30, 31^. We therefore treated four NC patient-derived cell lines, 10-15, PER-403, TC-797, and 14169^32^, with taz for 4 to 14 days, using U2OS cells as non-NC control. Taz treatment for 4-6 days resulted in squamous differentiation, evidenced by marked morphological changes, including flattening, enlargement, and spreading of cells, and expression of involucrin (IVL, or the epithelial differentiation marker, AE1/AE3) (**Fig. 1B**-C). Squamous differentiation correlated with loss of H3K27me3 (**Fig. 1D**). All four NC cell lines were very sensitive to taz with 10-day IC50 values in the low-to-mid nM range, compared to µM IC50 for the osteosarcoma cell line U2OS that does not harbor BRD4-NUT (**Fig. 1E**). NC cells completely arrested growth at single digit µM concentrations, whereas complete growth arrest was not achieved in U2OS cells in the dose range used (**Fig. 1E**). These findings indicate that H3K27 methylation by EZH2 is required to maintain growth through the blockade of differentiation of NC.

### Genes de-repressed by EZH2 inhibition are enriched with H3K27me3

We next examined transcriptional changes due to inhibition of EZH2 by taz. We performed rRNA-depleted RNA sequencing to comprehensively identify differentially expressed (DE) genes in response to taz. Given that the phenotypic responses to EZH2 inhibition occur over many days (**Fig. 1B**-E), we collected RNA from NC cells at multiple time points to identify the earliest time of robust transcriptional response. Analysis of RNAseq quality revealed high replicate reproducibility for all time points from PER-403 treated cells, although there was some batch effect in 10-15 treated cells for all time points (Supplementary **Fig.** S1A-B).

Of four time points (36h, 72h, 96h, 144h), the earliest large increase in global differential gene expression in response to taz occurred at 96h (**Fig. 2A**, Supplementary **Fig.** S1C-D). We therefore identified DE genes common to both NC cell lines after 96h of treatment. As predicted, based on the canonical function of EZH2 in repressing gene transcription, the majority of DE genes were upregulated (DE up) in taz-treated samples relative to DMSO treated controls (**Fig. 2B**). Moreover, nearly half of the DE up genes (452 of 956, 47%) in 10-15 cells were also DE up in PER-403 cells (**Fig. 2C**).

**Fig. 2.**
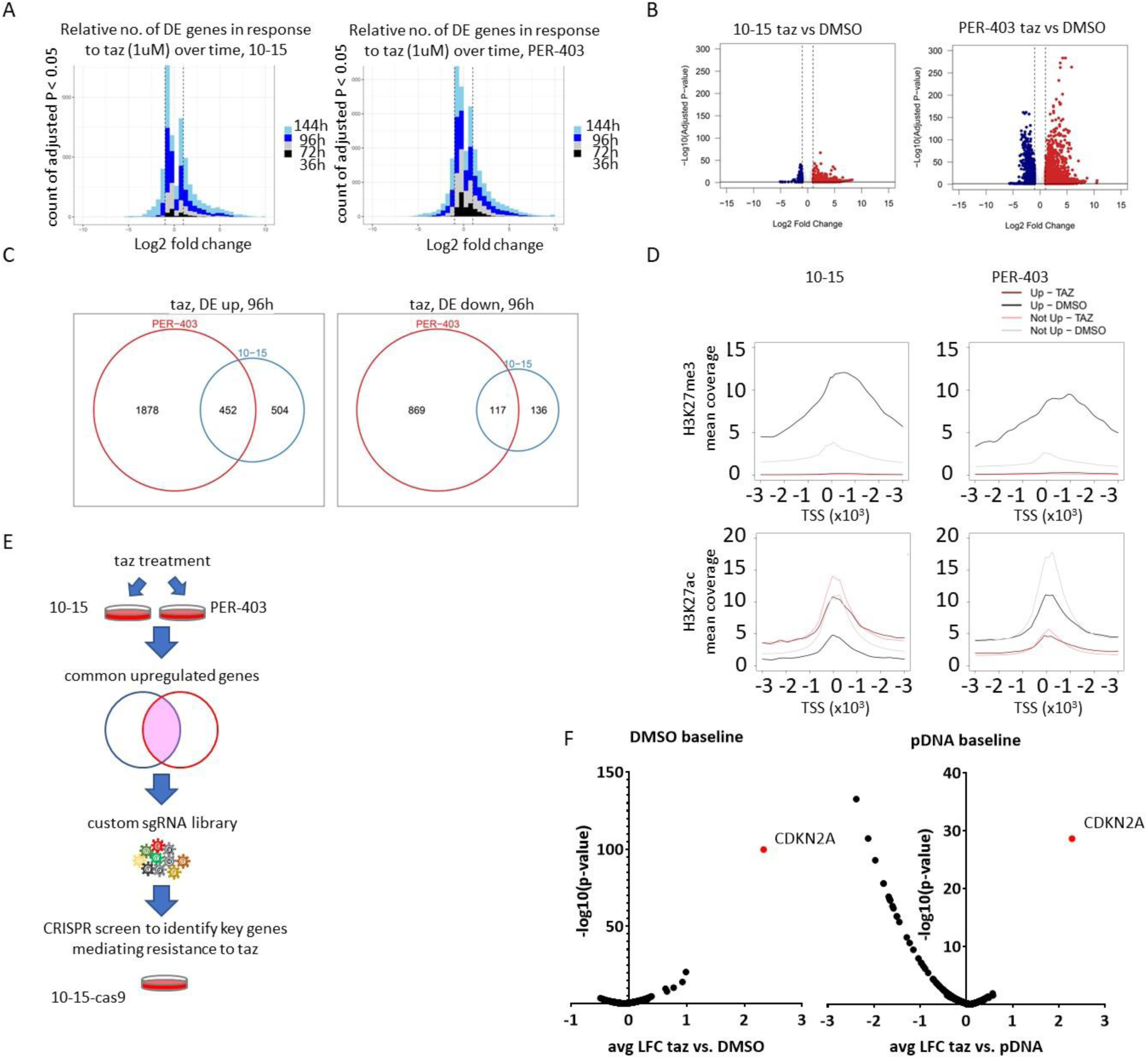
Genes repressed by EHZ2-induced H3K27 trimethylation include *CDKN2A*, a key NC tumor suppressor. A. Time series histograms of log2 FC for genes with adjusted p-value < 0.05 from RNAseq performed on samples from the indicated time points. B. Volcano plots depicting DE genes comparing RNAseq from DMSO-treated (72h) with taz-treated cells (96h). C. Venn diagrams of DE genes identified in taz treated samples. D. Enrichment profiles in H3K27me3- and H3K27ac-associated chromatin at the transcriptional start site (TSS) of coding genes. E. Schematic of strategy to identify key genes mediating resistance to taz treatment using a CRISPR-CAS9 screen. F. Plot of log (fold change) (LFC) averaged for each gene vs. p-values from the CRISPR-taz-resistance screen of 10-15-cas9 cells. Shown are representative single biological replicates from duplicate experiments. pDNA, plasmid DNA.

To reveal a mechanism underlying the changes in gene expression induced by EZH2i we examined repressive (H3K27me3) and active (H3K27ac) histone post-translational modification patterns genome-wide. We performed Cleavage Under Targets & Release Using Nuclease (CUT&RUN)^33^ for H3K27me3 and for H3K27ac with spike-in normalization on both 10-15 and PER-403 cells. As anticipated, taz treatment resulted in nearly complete, global eradication of H3K27me3.

The effects of taz on BRD4-NUT-associated H3K27ac domains, however, was mixed, depending on the cell line (Supplementary **Fig.** S1E-F). In 10-15 cells, there was an increase in global H3K27 acetylation, as predicted based on a previous study showing that EZH2i can induce compensatory increased H3K27ac in an MLL1-dependent manner^34^. In PER-403s cells, however, the opposite occurred, with marked decreased global H3K27ac (Supplementary **Fig.** S1E-F).

While the changes in global acetylation are intriguing, they are likely less functionally impactful than those from demethylation. It has recently been shown that H3K27 acetylation alone is not required for de-repression of genes silenced by EZH2-mediated H3K27me3^35^. Consistent with this observation, we found that genes upregulated by taz were more enriched for H3K27me3 at their transcription start sites (TSS) than those not upregulated by taz, in both DMSO-treated NC cell lines (**Fig. 2D**). As expected, H3K27me3 was completely depleted at genes in taz-treated samples regardless of their transcriptional response to taz. The opposite appeared to be true for H3K27ac, which was more highly enriched at the TSS of genes in DMSO-treated samples not up-regulated by taz than those upregulated by it, in both cell lines. Consistent with what was observed globally, H3K27ac increased at TSSs in taz-treated 10-15s but did the opposite in that of PER-403s (**Fig. 2D**). Overall, the CUT&RUN analysis indicates that gene de- repression by EHZ2 inhibition is due to loss of H3K27me3 in NC.

### *CDKN2A* is a key NC tumor suppressor gene directly repressed by H3K27me3 in a EZH2i- reversible manner

Having revealed that taz inhibition of EZH2-catalyzed H3K27me3 arrests NC cell growth and promotes differentiation, we sought to identify, in an unbiased manner, specific genes regulated by EZH2. Our overall approach is illustrated in **Fig. 2E**. We designed a custom CRISPR sgRNA library targeting the common DE up genes using four sgRNAs per gene (Supplementary Table S1). We also created a stable derivative of 10-15 cells that express Cas9 (Supplemental **Fig.** S1G-H). We transduced 10-15-Cas9 cells with our custom guide-only lentiviral sgRNA library, selected for transduced cells with puromycin for five days, and then treated cells with taz or vehicle (DMSO) for 14 days. This is a taz-resistance screen, where cells lacking key genes that confer a selective growth advantage in the presence of taz outcompete cells sensitive to taz. In this manner, genes whose repression mediate the pro-growth effects of EZH2 methyltransferase activity are identified. Quality analysis of the CRISPR screen indicated excellent replicate reproducibility of positively and negatively selected outliers comparing fold-change of taz- selected hits relative to the plasmid DNA library (pDNA), and likewise for positively selected outliers relative to DMSO-treated samples (Supplementary **Fig.** S1I).

Comparison of taz-selected genes with that of DMSO control or the pre-selected library revealed a single strong positive hit for all sgRNAs targeting *CDKN2A* (**Fig. 2F**, Supplementary **Fig.** S1J).

To test whether CDKN2A serves a tumor suppressor function to regulate growth of NC, we induced over-expression of the p16INK4a isoform of CDKN2A. p16INK4a induction, compared with GFP-induced negative control, led to near-complete blockade of growth corresponding with G1-cell cycle arrest and morphologic flattening of NC cells resembling senescence (**Fig. 3A**-D, Supplementary **Fig.** S2A). p16INK4a-induced growth arrest was associated with reduced phosphorylation of RB (**Fig. 3B**), a pivotal cell cycle regulator downstream of p16INK4a.

**Fig. 3.**
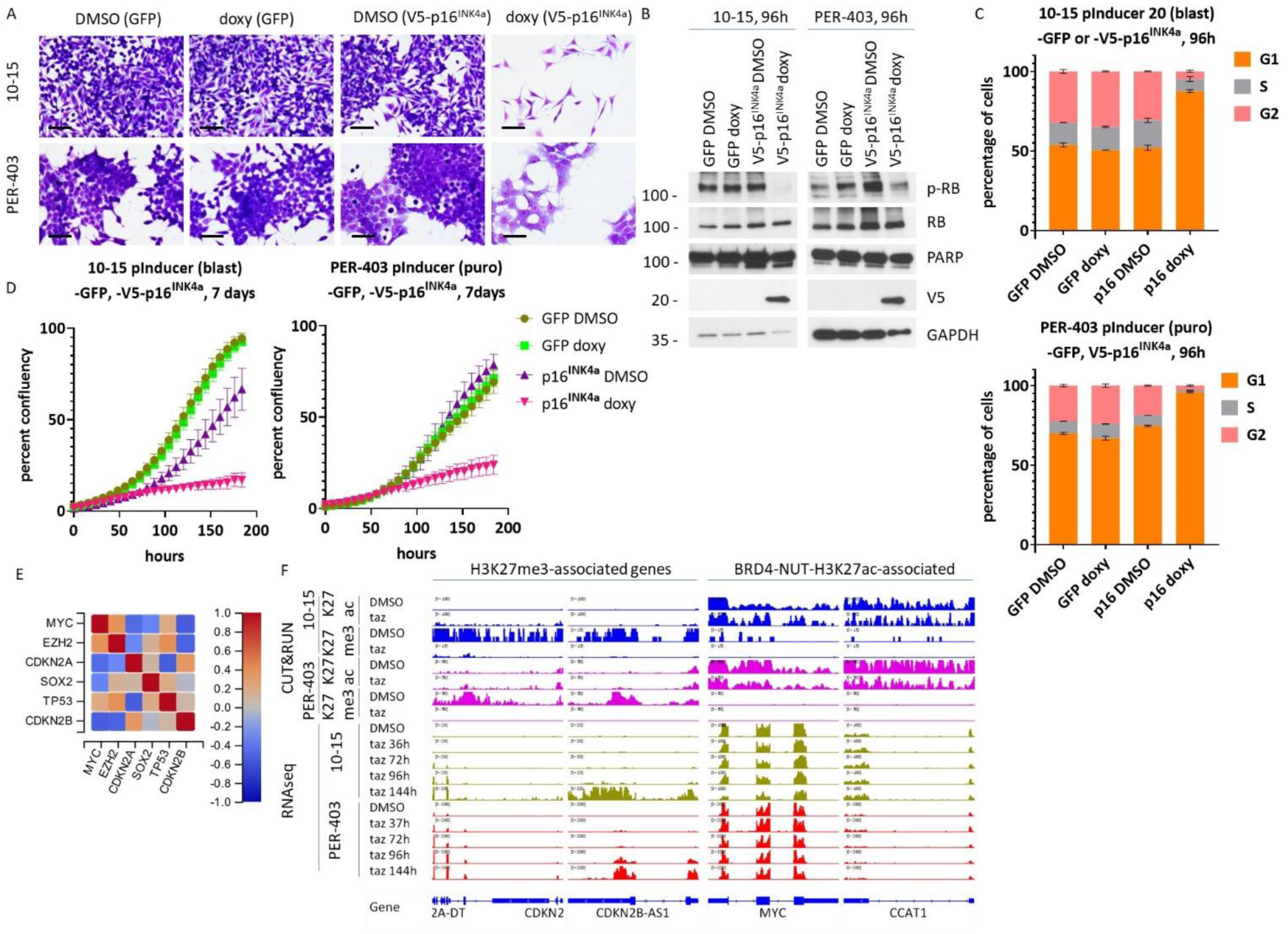
CDKN2A is a key tumor suppressor in NC. A. Hemacolor-stained NUT carcinoma cells transduced by pInducer 20 lentiviral plasmids induced to express the indicated cDNAs for 96 h. Scale bar, 25µm. B. Immunoblots corresponding with experiment in A. C. Flow cytometric analysis of NC cells treated as indicated. D. Cell confluency measured by incucyte imaging over seven days. E. Correlation plot derived from GeoMx DSP performed on the NC TMA. F. Integrated genome viewer views of CUT&RUN and RNAseq peaks at the indicated genes. Each track shown is from one of two biologic replicates. K27ac, CUT&RUN using anti-histone H3 K27ac; K27me3, CUT&RUN using anti- histone H3 K27me3.

To establish the *in vivo* role of CDKN2A in NC, we examined patient *CDKN2A* expression levels from the NC TMA. Expression of *CDKN2A* and *CDKN2B* profiled with the GeoMx^®^ Digital Spatial Profiler (DSP) strongly anti-correlated with that of *MYC* and *EZH2* (**Fig. 3E**). This correlative data suggests that EHZ2 may antagonize the expression of these two tumor suppressor genes. Taken together, the taz-resistance CRISPR-Cas9 screen and blockade of growth induced by p16INK4a strongly point to CDKN2A playing a key tumor suppressive function in NC.

Chromatin profiling by CUT&RUN corroborated the genetic evidence that *CDKN2A* confers sensitivity of NC cells to taz. Both *CDKN2A* and *CKDN2B*, amongst others, were highly enriched for H3K27me3, but not H3K27ac. Upon taz treatment, *CDKN2A* and *CKDN2B* were de-repressed, corresponding with complete erasure of H3K27me3 (**Fig. 3F**). By contrast, key BRD4-NUT-H3K27ac-associated genes, including *MYC* and *CCAT1*, were unaffected by taz treatment (**Fig. 3F**). This mechanistically demonstrates that direct repression of *CDKN2A* by H3K27me3 confers sensitivity of NC cells to taz.

### H3K27me3 and H3K27ac domains do not co-localize

We further examined the types of genes enriched for H3K27me3 by CUT&RUN. Multiple tumor suppressor genes, in addition to *CDKN2A* and *CKDN2B*, including *IGFBP3*, *GJB2*, *PLK2*, and *SOCS3* (Supplementary **Fig.** S2B), were highly enriched with H3K27me3, but not H3K27ac. These genes, classified as tumor suppressors in the TSGene database^36^, were also derepressed by inhibition of EZH2 with taz. This suggests that EHZ2 and BRD4-NUT may regulate separate chromatin domains, and therefore genes, in a complementary manner. To examine this in a more rigorous manner than bulk chromatin analysis by CUT&RUN, we performed confocal immunofluorescence (IF) imaging to correlate the spatial localization of H3K27me3 and H3K27ac domains in single cells. Co-localization of H3K27ac and BRD4-NUT is robust enough that H3K27ac can be used as a surrogate marker of BRD4-NUT for IF^8, 10, 14^. We began by establishing controls for ideal colocalization and negative colocalization. For ideal co- localization, we stained cells with anti-H3K27ac and co-detected using secondary antibodies from two different species. We then calculated the Pearson correlation between the two IF signals for one image each of 24 nuclei; this yielded a high correlation coefficient, thus establishing colocalization. We separately co-stained cells with antibodies to anti-H3K27ac and anti-NUP62, the latter a nuclear envelop- associated protein. These yielded a low Pearson correlation coefficient, thus establishing a lack of colocalization benchmark. H3K27ac-BRD4-NUT-associated domains are visualized as nuclear puncta^8, 10, 13^, whereas H3K27me3 is more diffusely distributed throughout the nucleus (**Fig. 4A**-B). We found that in both 10-15 and PER-403 NC cell lines, the correlation coefficient between H3K27me3 and H3K27ac domains trended towards the correlation coefficient between H3K27ac and NUP62, thus indicating a lack of colocalization (**Fig. 4C**). Taken together, the overall findings indicate that a large proportion of BRD4- NUT-H3K27ac chromatin domains are physically and functionally separate from EZH2-H3K27me3 domains. Such physical separation would explain why different genes are regulated by each epigenetic regulator.

**Fig. 4.**
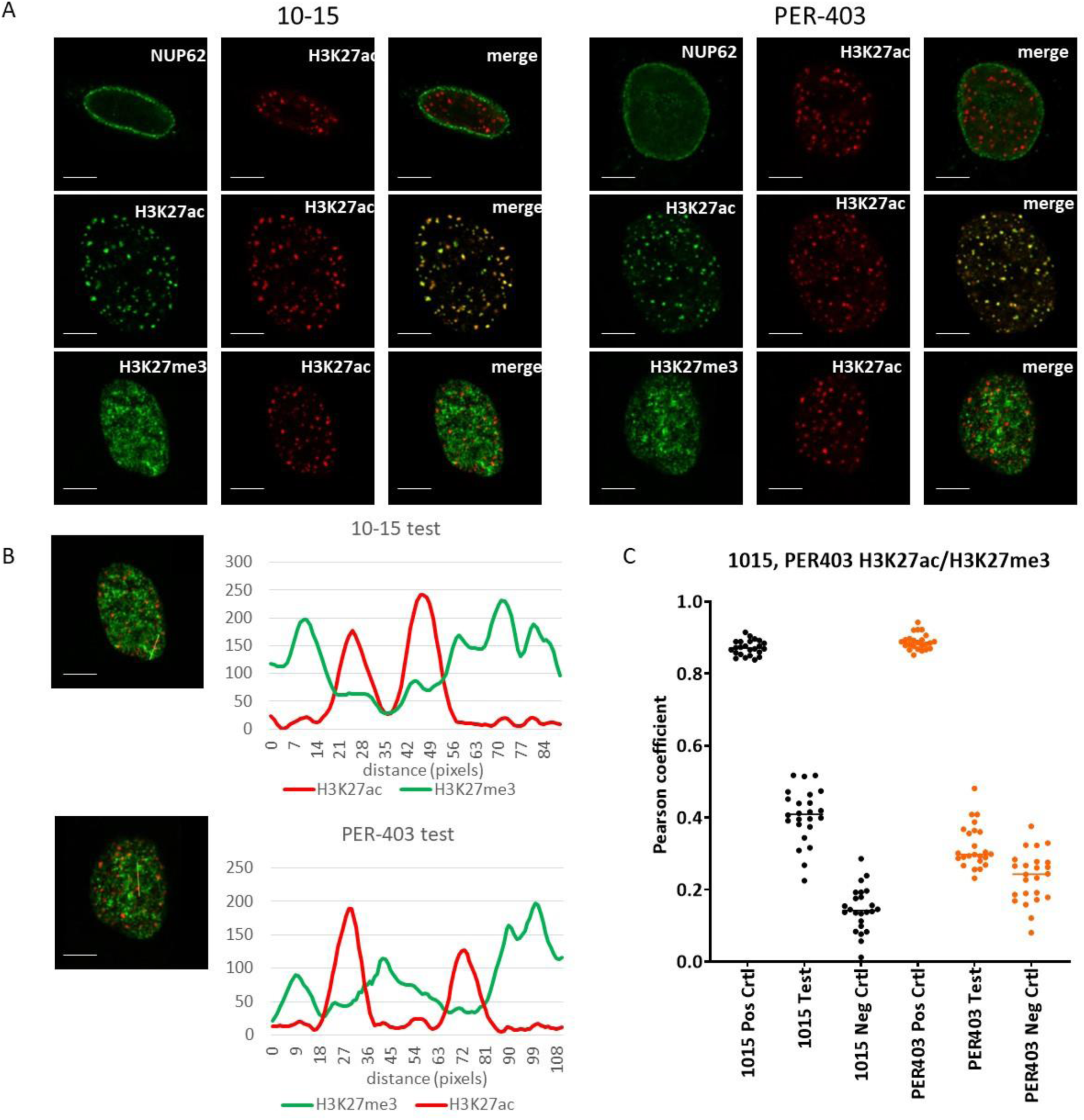
H3K27me3 and H3K27ac domains do not co-localize. A. Confocal immunofluorescent images as indicated. Scale bars, 3µm. B. Line-scan profiles of confocal immunofluorescent images to left (generated from A). Scale bar, 5µm. Profiles were generated using ImageJ. C. Pearson correlation of H3K27ac and H3k27me3 localization. 24 nuclei per group were analyzed.

### Combined EZH2 and BET inhibition synergistically induces terminal differentiation and blocks growth of NC

A physical separation of BRD4-NUT-H3K27ac and EZH2-H3K27me3 chromatin domains that contain functionally distinct sets of genes suggests that inhibiting both BRD4-NUT and EZH2 may be more effective in blocking NC growth than inhibiting either pathway alone. We therefore used taz to inhibit EZH2, and two BETi compounds, ABBV-075 and ABBV-744, to inhibit BRD4-NUT. ABBV-075 inhibits bromodomains 1 and 2 (BD1 and BD2) with roughly equal IC50s^37^. ABBV-744 is a BD2-selective BET inhibitor and shows clinical promise in a small subset of cancers, including prostate cancer, due to its much more favorable toxicity profile^38^. All five NC cell lines were exquisitely sensitive to both ABBV-075 and -744 *in vitro*, compared with the non-NC cell line, 293T (Supplementary **Fig.** S3A). Moreover, combined taz and BETi (both ABBV-075 and -744) synergistically inhibited NC growth at most dose combinations (**Fig. 5A**-B). The combination was highly synergistic in promoting squamous differentiation at very low concentrations of taz (80-100nM) and BETi (0.1-0.5 nM, **Fig. 5C**-D and Supplementary **Fig.** S3B-C). The synergistic anti-proliferative and pro-differentiation effect of combined taz and BET inhibition correlated with a greater proportion of G1-arrested cells compared to cells treated with either compound alone (**Fig. 5E**, Supplementary **Fig.** S3D).

**Fig. 5.**
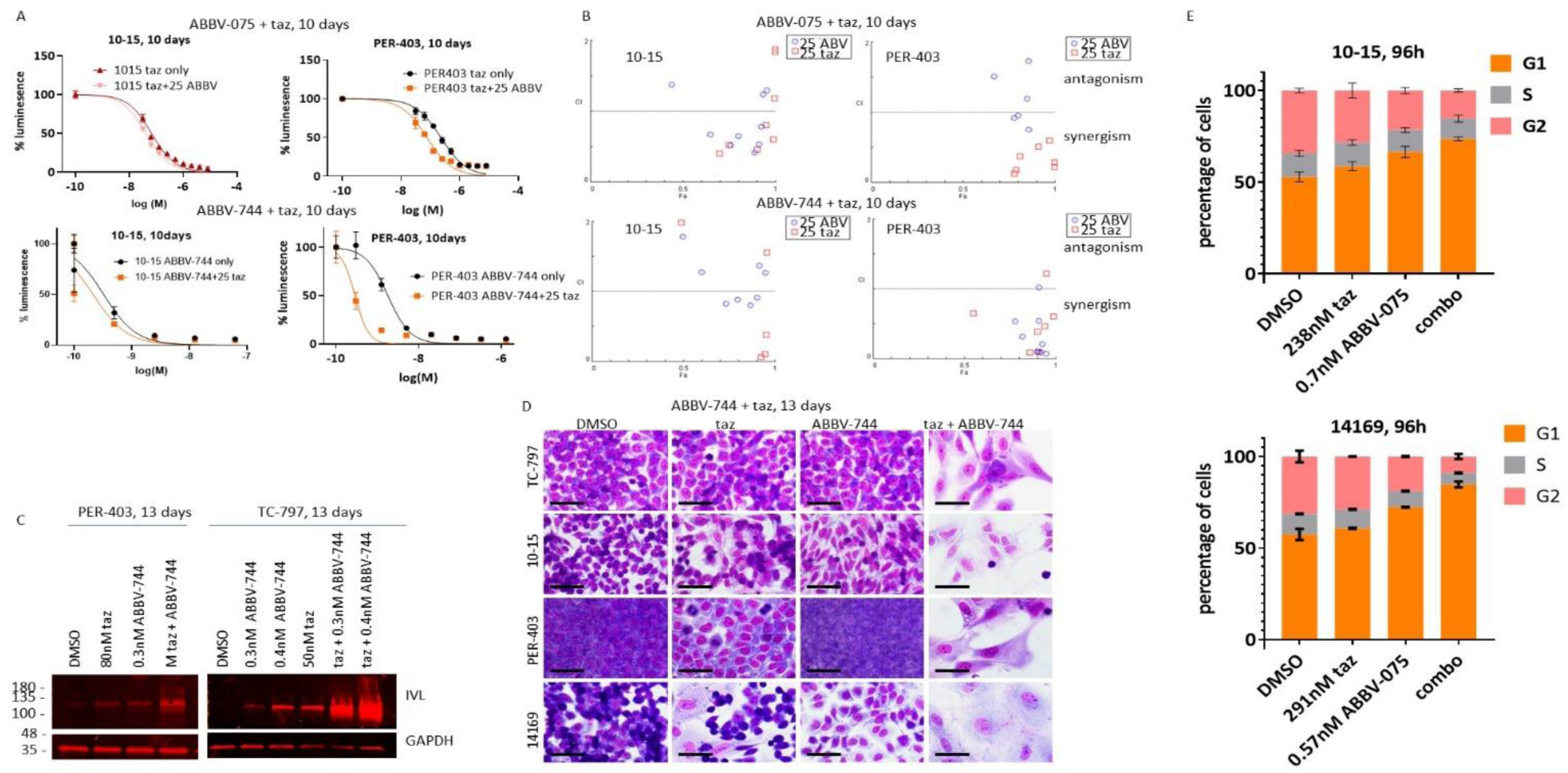
Combined EZH2 and BET inhibition synergistically induces terminal differentiation and blocks growth of NC. A. Dose-response to combinations of taz and ABBV-075 or ABBV-744. Top, variably dosed taz with or without fixed concentration of ABBV-075. taz + 25 ABBV, variable taz in combination with fixed dose of ABBV-075 (25% of IC50 for specified cell line). Bottom, variably dosed ABBV-744 with or without fixed concentration of taz. ABBV-744 + 25 taz, variable ABBV-744 in combination with fixed dose of taz (25% of IC50 for specified cell line). Dose response curves are from representative single replicates using Cell Titer Glo as readout. B. Chou-Talalay fraction affected (FA) vs combination index (CI) plots corresponding to treatments in A. 25 ABV, variable taz in combination with fixed (25% IC50) ABBV-075 or -744. 25 taz, variable ABBV-075 (or -744) in combination with fixed (25% IC50) taz. C. Immunoblots as indicated. D. Hemacolor stained cells grown on coverslips. Scale bar, 25µm. E. Flow cytometric analysis of NUT carcinoma cells treated as indicated.

### EZH2 and BRD4-NUT regulate separate sets of genes in NC, but combined inhibition synergizes to downregulate cell proliferation genes

To better understand the phenotypic synergy of combined EZH2i and BETi we analyzed corresponding changes in the transcriptional and epigenetic landscape in NC cells. Because BET inhibitors exert their effect within hours^8, 39^, we performed RNA-seq on 10-15 and PER-403 cells treated with ABBV-075 or ABV-744 after both short (6 hour) and long (96 hour) treatments. We treated with either BETi to determine if these two compounds have differing effects on gene expression, but found minimal-to-no differences in DE genes induced by ABBV-075 and -744 (Supplementary **Fig.** S4A). We also performed RNA-seq on 10-15 and PER-403 cells treated with taz for 96 hours. As before, the majority of DE genes were differentially upregulated in taz-treated samples relative to DMSO control, and taz-upregulated genes were more enriched for H3K27me3 at their TSSs than those not upregulated by taz (Supplementary **Fig.** S4B-C). We found that a substantial proportion of DE genes were unique to either BETi or EZH2i, with 40-59% non-overlapping DE genes. (**Fig. 6A**). This indicated that the expression of independent sets of genes were affected by the two inhibitors.

**Fig. 6.**
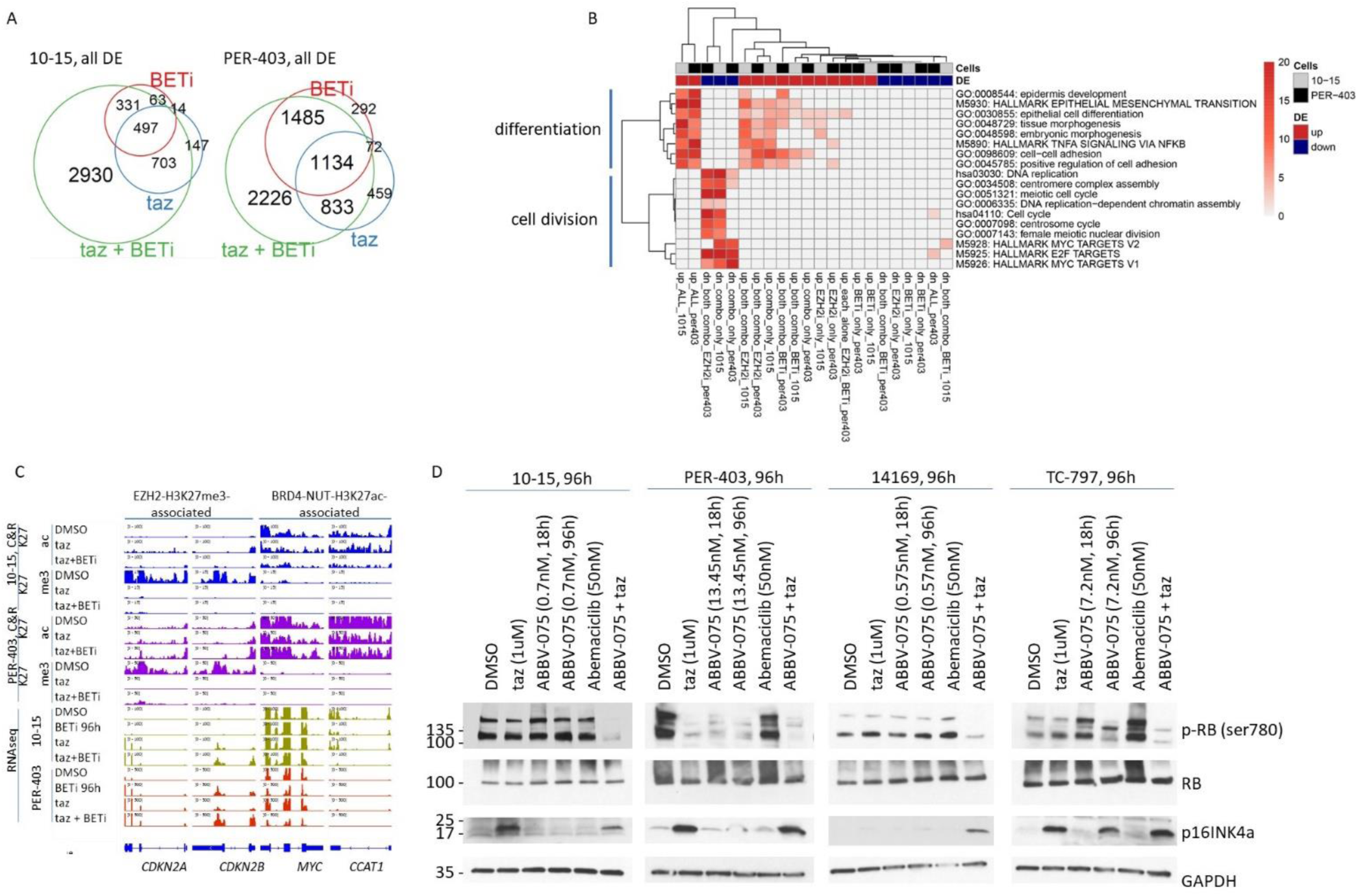
Combined BET and EZH2 inhibition synergizes to downregulate cell proliferation genes in association with loss of RB phosphorylation. A. Venn diagrams of DE genes identified by RNAseq analysis comparing cells treated by inhibitor with those treated with DMSO only for 96h. B. Metascape analysis comparing differentially up- and down-regulated genes for both 10-15 and PER-403 cell lines corresponding with data in A. C. Integrated genome viewer views of CUT&RUN and RNAseq peaks at the indicated genes. Each track shown is from one of two biologic replicates. D. Immunoblots as indicated.

We concurrently performed RNA-seq on 10-15 and PER-403 cells treated with taz combined with ABBV-075 for 96 hours. 34-62% of all DE genes were unique to the combined treatment (**Fig. 6A**). Metascape analysis of genes in the different areas of the Venn diagram revealed that while all compounds, alone or combined, appeared to induce epithelial-mesenchymal transition programs that induce epithelial/epidermal differentiation in both 10-15 and PER-403 cells, only the combination appeared to strongly inhibit pro-growth programs (MYC, E2F, meiotic, and cell cycle pathways, **Fig. 6B**- C, Supplementary **Fig.** S4D). Overall, the transcriptomic analysis closely matched the phenotypic findings of arrested growth and terminal differentiation induced synergistically by combined EZH2i and BETi.

### Combined EHZ2 and BET inhibition leads to synergistic loss of RB phosphorylation

Because CDKN2A is an important target of EZH2 in NC, we sought to determine the downstream effects of EZH2i, alone or combined with BETi. A central tumor suppressor function of the CDKN2A isoform, p16INK4a, is to activate RB by blocking its phosphorylation by CDK4/6. Thus, upregulation of CDKN2A expression by EZH2i is predicted to decrease levels of phospho-RB (p-RB) the same way that CDKN2A over-expression does (**Fig. 3B**). In fact, we found that while taz treatment increased expression of p16INK4a, only the combination of taz and ABBV-075 consistently depleted levels of p-RB, in four NC cell lines tested (**Fig. 6D**). Not even low doses of abemaciclib, a CDK4/6 inhibitor, affected p-RB. The marked decrease in p-RB seen in EZH2i + BETi-treated samples correlates strongly with the downregulation of cell division, E2F, and MYC transcriptional pathways exclusively by this combination or EZH2i (**Fig. 6B**). We hypothesize that the synergistic inhibition of RB phosphorylation may result from blockade of two key sources of RB phosphorylation: 1. inhibition of CDK4/6 by EZH2i via CDKN2A, and 2. transcriptional downregulation of CDK2/CDC25A^40^ by combined EZH2i and BETi, the latter evidenced by RNAseq (Supplementary **Fig.** S4D).

### Combined EZH2 and BET inhibition synergistically blocks NC *in vivo*

The *in vitro* potency of combined EZH2i and BETi motivated us to evaluate the activity of this combination in an *in vivo* pre-clinical model of NC. To model NC, we inject, via the tail vein, NOD-SCID-GAMMA (NSG) mice with 10-15, PER-403, or 14169 BRD4-NUT+ cell lines that express luciferase^39^. Upon injection these cells disseminate to solid organs, such as ovary and liver, and bone, within one-to-two weeks, robustly recapitulating the human disease. After disseminated tumor is established by bioluminescence imaging (BLI), we administered vehicle control, BETi, EZH2i, or combined BETi and EZH2i for 28 days, after which treatment is terminated and mice are observed for up to 170 days. Mice are treated with ABBV-744 (1/2 maximal tolerated dose (MTD), 37.5mg/kg qd), taz (250mg/kg BID), or ABBV-744 + taz.

In the PER-403 xenograft model (n = 4 mice per arm), there was no effect on growth nor survival benefit in mice treated with taz alone [p = 0.9452 (Log-rank (Mantel-Cox) test]. While there was an overall survival benefit in mice treated with ABBV-744 alone [p = 0.0067], all tumors grew continuously in the presence of single agent, and all mice treated as such were dead within 63 days (**Fig. 7A**). By contrast, all tumors treated with the combination regressed to baseline, and all mice were cured of disease as of 169 days follow-up, showing significant survival benefit compared with ABBV-744 alone (p = 0.0067). The effect of this combination on tumor growth and mouse survival is unprecedented in these (or any) *in vivo* models of NC, to our knowledge^17, 39, 41–44^.

**Fig. 7.**
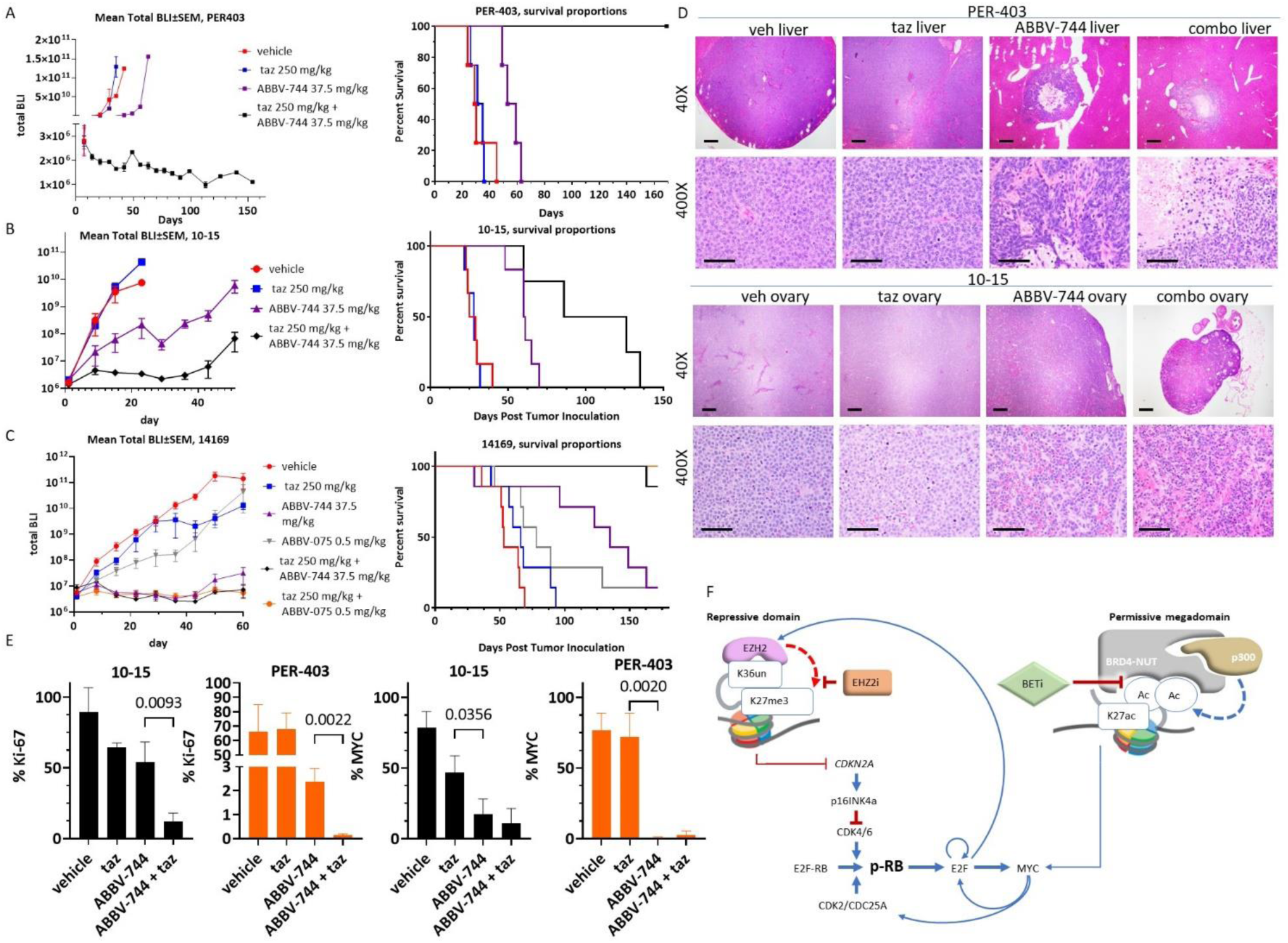
Combined EZH2 and BET inhibition synergistically blocks NC *in vivo*. A-C, left, tumor growth over time measured by total bioluminescence (BLI). Treatment is initiated on day 1, and is continuously administered until treatment termination on day 28. Right, Kaplan-Meier survival plots corresponding to left. D. Histologic and immunohistochemical analysis of tumors harvested from mice treated for 5 days with compounds indicated. Scale bar (40x images), 500µm; 400x images, 50µm. E. Ki-67 proliferation indices determined by positively stained cells counted in tumors harvested from three mice per treatment. 400x images were analyzed in ImageJ for cell counting. F. Cartoon of proposed mechanism of taz + BETi synergy.

In the 10-15 xenograft model (n = 6 mice per arm) we observed a similar overall survival benefit and repression of tumor growth using ABBV-744 as a single agent compared with taz or vehicle (p = 0.0005). Moreover, there was greater repression of tumor growth and significantly improved overall survival in mice treated with combined taz and ABBV-744 compared with monotherapy with ABBV-744 (p = 0.0333), however all mice eventually succumbed to disease progression by day 135 (**Fig. 7B**). While no cures were seen, the improved survival resulting from this combination is also unprecedented for this particularly resilient cell line xenograft model^39^.

In the 14169 model (n = 7 mice per arm), we additionally tested the efficacy of the pan-BET bromodomain inhibitor, ABBV-075, alone or combined with taz. As for ABBV-744, we dosed ABBV-075 at half the maximal tolerated dose (0.5 mg/kg). The efficacy of both combinations resembled that seen in the PER-403 model, with only one death in seven mice observed in the ABBV-744 + taz group, and zero deaths in four mice seen in the ABBV-075 + taz group, by the end of the study on day 170 (**Fig. 7C**). Both combinations provided significantly improved overall survival compared with either BETi alone (ABBV-744 vs ABBV-744 + taz: p = 0.0039; ABBV-075 vs ABBV-075 + taz: p = 0.0139; ABBV-075 vs ABBV-744 + taz: p = 0.0005). By study end, three mice in the ABBV-075 + taz, and two in the ABBV-744 + taz had minimal-to-no detectable tumor [1.7e6 – 8.4e6 photons/sec/cm^2^/steradian (p/s/cm^2^/sr)], whereas the remaining mice had moderate-to-high tumor burden (5.8e8 – 7.96e11 p/s/cm^2^/sr). The effects on overall survival are again unprecedented in any xenograft model of NC, to our knowledge^17, 39, 41–44^.

Pharmacodynamic evaluation of two xenograft models, 10-15 and PER-403, was performed on tumors removed from mice comparing five-day treatment with ABBV-744 (37.5 mg/kg), taz (250 mg/kg), and ABBV-744 + taz, with vehicle. Histologically, tumors were markedly smaller in ABBV-744 and ABBV- 744 + taz-treated tumors, becoming almost undetectable in the combination group; the small tumor sizes in the latter two groups corresponded with the presence of extensive hemorrhage, necrosis (**Fig. 7D**). Proliferation, scored by Ki-67 proliferation index, was significantly decreased in ABBV-744-only-treated animals compared with taz-only, and in ABBV-744 + taz-treated compared with ABBV-744-only-treated mice in both models (**Fig. 7E**, Supplementary **Fig.** S5A). Likewise, MYC expression was significantly reduced in ABBV-744- and ABBV-744 + taz-treated mice compared with taz-only (**Fig. 7E**, Supplementary **Fig.** S5A). As predicted, H3K27me3 was decreased in taz-treated tumors, and this corresponded with increased p16INK4a expression (Supplementary **Fig.** S5B).

Taken together, the pre-clinical activity of combined EZH2i and BETi is highly synergistic in our three models, and far surpasses that observed in any NC pre-clinical model to date. Immunohistochemical data indicates that the drugs act *in vivo* as expected based on our *in vitro* data. This data provides strong rationale for clinical evaluation of combined EZH2i and BETi in patients with NC.

## DISCUSSION

The identification of EZH2 as a critical dependency in NC is notable for two reasons: (1) substantiation of NC as an epigenetic disease of aberrant gene expression, and (2) development of a synergistic drug combination with unprecedented *in vivo* activity. Silencing of tumor suppressor genes by EZH2 is significant as it identifies a pathway parallel to BRD4-NUT-induced gene activation to maintain NC cell viability. Importantly, both pathways are epigenetic, yet have opposing outcomes on gene activity. Simultaneous pharmacologic targeting of complimentary epigenetic pathways led to drug synergy; combined EZH2 inhibition and BET inhibition was more efficacious than either inhibitor alone.

Our study shows that EZH2 complements BRD4-NUT by repressing at least one key tumor suppressor gene, *CDKN2A*, in NC. *CDKN2A* encodes two proteins, p14ARF and p16INK4a that function to regulate cell cycle progression, and in some circumstances, induce senescence. p14ARF functions to inhibit degradation of p53 by MDM2^45^, and p16INK4a inhibits inactivation of RB by CDK4/6^46^. CDKN2A is commonly inactivated by a variety of mechanisms in cancer, most often by deletion^47^, DNA promoter methylation^48^, or mutation^48^. In cancers where *CDKN2A* is not inactivated by mutation or promoter methylation, it must be repressed by other mechanisms, often epigenetically by PRC2^49^. NC appears to be one of those cancers.

*CDKN2A* is not the only tumor suppressor gene repressed by EZH2 in NC. We find several other tumor suppressor genes down-regulated by EZH2, including *CDKN2B*, *IGFBP3*^50, 51^, *GJB2*^36^, *PLK2*^52^, and *SOCS3*^53^ are highly tri-methylated at H3K27 and de-repressed by EZH2i (**Fig. 3F**, Supplementary **Fig.** S2B). p15INK4b, which is encoded by *CDKN2B*, serves a similar function as p16INK4a to inhibit CK4/6 phosphorylation of RB. IGFBP3 is activated by p53 to block growth and induce apoptosis^50, 51^, PLK2 may block tumor growth through regulation of PLK1^52^, and SOCS3 may suppress tumor growth through repression of STAT signaling^53^ . These genes were included in our CRISPR screen, but their loss failed to provide a selective growth advantage to NC cells in the presence of taz. Possible reasons include depletion negatively affecting fitness in the presence of taz or alternatively, non-tumor suppressor functions. Regardless, the oncogenic function of EHZ2 in NC may extend beyond simply repressing *CKDN2A*.

One of the most notable findings in this study is the profound synergy resulting from combined EZH2i and BETi, manifested as terminal squamous differentiation and arrested growth, both *in vitro* and *in vivo* (**Fig. 5**, 7, Supplementary **Fig.** S3, S5). The complementary, yet distinct functions of BRD4-NUT and EZH2-associated pathways provides an explanation for this synergy. Synergy of this combination has also been reported in another study of aberrant EZH2-expressing non-NC tumors. Huang et al^34^ found that, mechanistically, BETi counteracts compensatory MLL1-mediated global acetylation of H3K27 in response to H3K27me3 demethylation by EZH2i^34^. We observed both global increases and decreases in H3K27 acetylation in our cell lines in response to taz (Supplementary **Fig.** S1E-F), thus the synergy we observed in NC cannot occur through this mechanism. Rather, we consistently observed unambiguous downregulation of cell cycle and cell proliferation pathways related to MYC and E2F targets in both NC cell lines tested (**Fig. 6B**).

We propose a mechanistic model of how EZH2i and BETi synergistically block growth of NC (**Fig. 7F**). The model is consistent with our previous demonstration that *MYC* is a direct target of BRD4-NUT, and that BETi reduces MYC levels in NC^8, 9^. Likewise, it is established that functional E2F requires phosphorylation of RB by CDK2/4/6; thus de-repression of CDN2A by EZH2i is expected to block transcriptional upregulation of S-phase genes by E2F. *MYC* is a known transcriptional target of E2F as well^54^, thus reduced E2F by EZH2i is expected to cooperate with BETi to lower MYC levels further, which we observed by RNA-seq (**Fig. 6C**, Supplementary **Fig.** S4D). Furthering this feed-forward cycle is that *EZH2* is transcriptionally upregulated by E2F^55^. This results in feed-forward regulation such that E2F’s inhibition by CDKN2A is expected to reduce transcription of EZH2, which is also seen by RNAseq (Supplementary **Fig.** S4D). Lastly, we observed that there was a combinatorial effect of EHZ2i + BETi on RB de-phosphorylation. This possibly occurs due to a cooperative effect of reducing expression of CDK2 and its activating phosphatase, CDC25A, a transcriptional target of MYC^56^ (Supplementary **Fig.** S4D). Together this increases p-RB. The profound effect on RB de-phosphorylation likely explains the strong reduction of E2F-associated transcriptional pathways (**Fig. 5B**). This effect is further amplified by the reduced expression of E2F1 itself, a transcriptional target of MYC^57^(Supplementary **Fig.** S4D). In summary, central to our model is that the synergistic inhibition of cell proliferation by EZH2i + BETi stems from the multi-pronged de-repression of RB, and inhibition of MYC (**Fig. 7F**).

Notably, taz alone is completely ineffective *in vivo*, highlighting the synergy that occurs when it is combined with BETi. EZH2i is likely ineffective as a single agent because it poorly inhibits expression of *MYC* (**Fig. 3F**, 6B), the central driver of NC tumor growth; but also possibly because of its slow kinetics - delayed activity of taz monotherapy may be too slow to “catch up” with the exponential tumor growth of NC. EZH2 has recently been shown to non-canonically bind MYC and recruit p300 to co-activate MYC target genes^58^. EZH2 can stabilize MYC^59^; thus, degradation of EHZ2 using targeting PROteolysis-TArgeting Chimeras (PROTACs) is significantly more potent than catalytic EZH2 inhibitors^58^. EZH2 degradation may therefore phenocopy the effects of MYC inhibition when EZH2i is combined with BETi. *in vivo*-optimized EZH2 degraders would therefore have therapeutic promise in NUT carcinoma and other cancers.

NC was previously shown to be co-dependent on BET activity and RB loss/cyclin D1 gain, consistent with the results here^42^. CDK4/6 inhibition (CDK4/6i) by palbociclib exploited this vulnerability and synergized with BETi^42^. However, our *in vivo* EZH2i + BETi treatment is more durable than that reported for CDK4/6i + BETi^42^. When comparing CDK4/6i + BETi to EZH2i + BETi, additional factors should also be considered, such as different BET inhibitors used in the two studies or different treatment regimens. Regardless, our treatment regimen with EZH2i + BETi prevented tumor growth, even after treatment cessation, whereas tumor grew rapidly once CDK4/6i + BETi was withdrawn^42^. Additionally, EZH2i + BETi treated mice in our study survived over three times longer than those treated with CDK4/6i + BETi treatment^42^. These differences likely highlight that NC is an epigenetic disease. Inhibiting two parallel, complimentary epigenetic pathways is more effective that inhibiting parallel, but mechanistically distinct, epigenetic and signaling pathways. As shown, the global loss of H3K27me3 by EHZ2i (Supplementary **Fig.** S1E) derepresses 100s-1000s of genes (**Fig. 2B**-C), including the aforementioned numerous tumor suppressor genes. These changes likely add to the robustness and durability of the *in vivo* responses of NC cells to EZH2i + BETi compared with CDK4/6i + BETi. Pharmacokinetics also likely plays a role. For example, reestablishment of H3K27me3 after EZH2i is slow, taking 4-7 days^28, 29^ whereas RB phosphorylation by CDK4/6 can occur within 9h after restoration of cyclin D1 expression^60^.

A significant finding of our study is the mechanism by which NC is co-dependent on EZH2 and BRD4- NUT: RB inactivation by EHZ2 on the one hand, and MYC activation by BRD4-NUT on the other. It logically follows that combined inhibition of these two pathways is profoundly synergistic *in vivo*, which we demonstrate. This provides an opportunity to significantly improve therapy of this nearly uniformly fatal disease. Our work provides strong rationale for immediate clinical evaluation of combined EZH2i and BETi in NC, and other cancers that are co-dependent on these pathways.

## METHODS

### Tumor cell lines

NMC cell lines, TC-797^61^, 10–15^9^, PER-403^32^, 14169^62^, and non-NMC cell lines, 293T and U2OS, were maintained as monolayer cultures in DMEM (Invitrogen) supplemented with 10% FBS (Sigma-Aldrich batch number 17H392, St. Louis, MO) and 1% pen–strep (GIBCO/Invitrogen). The TC-797, PER-403, 14169, and 10–15 cell lines were authenticated by fluorescent *in situ* hybridization as described^16^ demonstrating rearrangement of the *NUTM1* and *BRD4* genes. The 293T and U2OS cell lines have not been authenticated.

### NUT carcinoma tumor micro-array (TMA)

Through our IRB-approved, NUT carcinoma international registry (www.NMCRegistry.org), we collected more than 100 formalin-fixed paraffin-embedded (FFPE) tissue blocks of primary NC. Select blocks of well-preserved tumor tissue (n = 77 different tumors) were used to create a TMA arraying 4 x 1mm cores from each (two cores from tumor margin and two cores from tumor center, 278 cores total) at the Dana Farber Harvard Cancer Center Pathology core, Longwood (Boston, MA).

### Immunohistochemistry

Formalin-fixed, paraffin-embedded tumor samples from mice were prepared using standard methods^15^. All immunohistochemistry (IHC) was performed on the Leica Bond III automated staining platform. Antibodies and conditions used are listed in Supplementary Table S2.

### Hemacolor cytologic staining

Adherent cells were grown on 25mm glass coverslips (Fisherbrand, Fisher Scientific, Pittsburgh, PA) in 6-well format, and stained using the MilliporeSigma Hemacolor Staining Kit for Microscopy according to the manufacturer’s instructions (MilliporeSigma, Temecula, CA).

### Histone extraction for immunoblotting

To evaluate levels of histone H3 histones in the chromatin, histone was extracted using a Histone Extraction Kit (ab113476, Abcam) according to the manufacturer’s instructions and analyzed by immunoblot.

### Immunoblotting

Cells were lysed in RIPA Buffer (50 mM Tris-HCl, 250 mM NaCl, 1% NP-40, 0.5% Sodium Deoxycholate, 0.1% SDS, 5 mM EDTA) containing 250 mM NaCl and Halt Protease and Phosphatase Inhibitor Cocktail (78445, Thermo Fisher Scientific) for 30 min at 4°C with rotation. The lysate was centrifuged at 16,100 x g, and the supernatant was collected. Immunoblotting was performed as described previously^43^. Antibodies and conditions used are listed in Supplementary Table S3.

### Plasmids, cloning, and viral transduction

Cas9: plenti-EF1a-Cas9-2A-blast and pXPR047 (Cas9-GFP reporter, Addgene# 107145) plasmids were provided by Dr. Kimberly Stegmaier. pXPR_003 sgRNA guide-only vector was provided by Dr. David Root. pInducer20 blast (Addgene) was purchased from Addgene (plasmid # 109334, Watertown, MA). pInducer20 puro was a gift from Dr. Stephen Elledge.

pInducer20 puro (or blast) -V5-p16INK4a and -eGFP lentiviral plasmids were created by gateway cloning gBlocks (synthesized by Integrated DNA Technologies, Inc. (IDT), Coralville, IA) of their coding sequences into pInducer20 puro (or blast) using Gateway LR Clonase II Enzyme Mix according to the manufacturer’s instructions (Thermo Fisher Scientific, Waltham, MA).

To create lentiviruses, 293T cells were transfected with with vector DNA, pRev, pTat, pHIV Gag/pol, and pVSVG by transfection using Lipofectamine 2000 (Invitrogen, Waltham, MA). Virus was collected 48h after transfection and purified using a 45 µm pore filter. Polybrene (4 µg/ml, Sigma) was used to transduce 10-15 or PER-403 cells, and the infected cells were subsequently selected with either puromycin (1 µg/ml; Sigma-Aldrich (St. Louis, MO) or blasticidin (Gibco, Billings, MT) for at least one week.

### Cell growth assays (Cell Titer Glo)

Cells were seeded into 96-well plates at a density of 500-1,500 cells per well in a total volume of 100 µl media. Compounds were delivered to triplicate wells using a HP D300e digital dispenser (Hewlett Packard, Spring, TX) at the ICCB-Longwood Screening Facility at Harvard Medical School (Boston, MA).

For long term (>96h) treatments, cells were re-fed every four days using an Agilent Bravo liquid handler (Agilent Technologies, Santa Clara, CA). Following a 72-96h or 10 day incubation at 37 °C, cells were lysed, and wells were assessed for total ATP content using a commercial proliferation assay (Cell TiterGlo; Promega Madison, WI). Three biologic replicates were performed. Estimates of IC50 were calculated by logistic regression (GraphPad Prism, Dotmatics, Boston, MA).

### Cell growth assays (IncuCyte)

10-15 and PER-403s cells stably transduced with pInducer lentiviral plasmids were seeded at 1000 cells per well in a 96 well plate, on day zero, and treatment with doxycycline (0.1ug/ml) or DMSO was initiated on day one. Treatments were performed in technical sextuplets. Plates were incubated on an IncuCyte S3 imager at the Human Neuron Core at Boston Children’s Hospital. Images to measure confluency were taken every 8 hours with five images per well. Treatment and imagining proceeded for 7 days. Exported data was analyzed in GraphPad Prism.

### Chemicals and compounds

EPZ-6438 (tazemetostat) was generously provided by Epizyme (now Ipsen Bioscience, Cambridge, MA). ABBV-744 and ABBV-075 were kindly provided by AbbVie, Inc. (Chicago, IL). Abemaciclib was purchased from MedChemExpress (Monmouth Junction, NJ). Dimethyl sulfoxide (DMSO) was purchased from Sigma-Aldrich.

### RNA extraction and library preparation for RNAseq

Whole RNA was extracted from live cultured 10-15 and PER-403 cells treated in biologic duplicates with the indicated concentrations of vehicle (DMSO), ABBV-744, ABBV-075, taz, or taz combined with ABBV- 075 using the RNeasy Plus kit (Qiagen). 1 ug of total RNA was used for the construction of ribosome-depleted sequencing libraries using the KAPA RNA HyperPrep Kit with RiboErase kit (HMR, Roche) according to the manufacturers’ instructions. ERCC ExFold RNA spike-in controls (Invitrogen; 4456739) were added according to manufacturer’s instructions to 1 µg of purified RNA from each sample to help normalize expression. Ribosomal RNA (rRNA) was depleted using the NEBNext rRNA Depletion Kit (NEB; E6350L) according to manufacturer’s instructions. Libraries were 50bp paired-end sequenced on a novaseq 6000 sequencer at the University of Chicago Genomics Facility (Chicago, IL, http://genomics.bsd.uchicago.edu/).

### Analysis of RNAseq

Reads were trimmed and verified for quality with Trim Galore! (https://www.bioinformatics.babraham.ac.uk/projects/trim_galore/). Trimmed reads were aligned with STAR^63^ to the human genome (GRCh38 primary assembly) and to the External RNA Control Consortium (ERCC9) spike-in sequences combined using default settings for gene counting (--quantMode GeneCounts) with Ensembl annotations (release 105, http://dec2021.archive.ensembl.org/Homo_sapiens/Info/Index) supplemented with ERCC entries. The range of reads mapped for the 10-15 TAZ time series was unusually large (8.6-fold) so aligned reads were downsampled to match the sample with the lowest number of uniquely aligned reads (TAZ t=144h). Transcripts with < 0.5 million reads mapped in all samples (n=2 per group) were removed before differential expression (DE) analysis. The R package RUVSeq^64^ was used to capture unwanted variation. RUVg was run with k=1 and invariant gene sets comprising either ERCC spike-ins or, for the TAZ time series (no spike-ins), genes not DE (adjusted p-value > 0.5) at any time vs the t=72 DMSO control. The resulting RUVg weight matrix (W_1) was incorporated into the DESeq2^65^ design matrix. P-value distributions were checked and found to have an anti-conservative Uniform distribution, as expected^66^. Genes were deemed to be differentially expressed with Benjamini-Hochberg^67^ adjusted p-value < 0.05 and a fold change (FC) > 2. Time series histograms of log2 FC in TAZ-treated samples vs t=72h DMSO controls were restricted to genes with adjusted p-value < 0.05. Venn diagrams were generated using R packages Vennerable (proportional Venn^68^) or gplots (petal plots^69^). Metascape analysis was run online (www.metascape.org) comparing differentially up- and down-regulated genes for both 10-15 and PER-403 cell lines with a background gene set consisting of genes with at least 0.5 mapped reads per million in at least one sample (Supplementary Table S4). Enrichment results were plotted in R using the pheatmap^70^ package with the -logP values clipped at 20 to increase color contrast.

### Analysis of CUT&RUN sequencing

Reads were trimmed for adaptor sequences using trimmomatic v0.36 and then aligned to concatenated hg38/Genome Reference Consortium Human Reference 38 (GRCh38) and dm6/BDGP Release 6 + ISO1 MT assemblies of the human and *Drosophila melanogaster* genomes, respectively, using bowtie2 v2.3.1 with options --local --very-sensitive-local --no-unal --nomixed --no-discordant --phred33. Only reads with a MAPQ ≥ 30 were retained using samtools v1.3.1. PCR duplicates were removed with picard MarkDuplicates v2.9.2. Coverage-based tracks were generated with deepTools bamCoverage with options --binSize 1 -normalizingUsing RPGC --effectiveGenomeSize 2800000000 -extendReads -- samFlagInclude 64. CUT&RUN profiles were then calibrated by spike-in normalization using the Drosophila genome as the internal reference using a similar method as that done for ChIP-seq^71^. Specifically, the Occupancy Ratio (OR) was calculated as the product of IgG spike-in reads and immunocleaved (IC) human reads, divided by the product of IgG human reads and IC spike-in reads. Spike-in normalized tracks were generated with deepTools bamCoverage with options --binSize 1 -- normalizeUsing RPGC --effectiveGenomeSize 2800000000 --extendReads --samFlagInclude 64 -- scaleFactor OR. epic2^72^ v0.0.52 was used for domain calling on the canonical human chromosomes with options --bin-size 200 --gaps-allowed 5 with the corresponding IgG library set as the control. Called domains were filtered by signal strength and by domain size using the ROSE algorithm (Rank Ordering of Super-Enhancers)^73^. Domains that exceeded both the signal cutoff and the size threshold were retained. We frequently observed closely spaced filtered domains that corresponded to larger domains based on CUT&RUN signal tracks. To span and capture these regions filtered domains within 25 kb of one another were combined. A high confidence list of “megadomains” was created by merging filtered domains identified in both biological replicates.

### CRISPR-Cas9 screen reagents and procedures

A custom lentiviral sgRNA library was created targeting differentially upregulated genes in response to taz that are common to 10-15 and PER-403 cell lines, using 4 sgRNAs per gene. We used the list of DE up genes common to PER-403 and 10-15 cells (Supplementary Table S1) to create a <u>custom lentiviral</u> <u>guide-only library for the taz-resistance CRISPR-Cas9 screen</u> using CRISPick (https://portals.broadinstitute.org/gppx/crispick/public. For this screen, we created a stable, blasticidin-selectable, Cas9-expressing derivative of 10-15 (Supplemental **Fig.** S2E-F). 26 common essential genes were included as positive controls, and 100 intergenic and 50 non-targeting guides were included as negative controls. The list of genes and 2046 total guide sequences is in Supplementary Table S1. The DNA oligonucleotide library was synthesized on a microarray (Agilent), and each gRNA was PCR amplified, digested with BbsI, and cloned into BsmBI-digested pXPR_003, at the Broad Institute Genetic Perturbation Platform as described^74^.

The CRISPR screen was conducted in duplicate. 10-15-Cas9 cells were infected with pooled CRISPR library virus using polybrene (8 µg/ml, Sigma) using a low multiplicity of infection (MOI = 0.75) to achieve an average representation of ∼500 cells per gRNA. The transduced cells were selected using puromycin (1 μg/ml; Clontech) for five days. After harvesting a subset of cells on day zero, the remaining cells were divided into two groups and treated with DMSO or 1µM taz for 14 days, and were then harvested. Bar- coded plasmid DNA was amplified via PCR and sent for next-generation sequencing (NGS) at the Broad Institute. Quantification of NGS data was performed using PoolQ (https://portals.broadinstitute.org/gpp/public/software/poolq), and STARS_v1.3 (https://portals.broadinstitute.org/gpp/public/software/stars) was used to rank the performance of individual genes based on enrichment comparing the taz treatment group with the DMSO-treated group or library plasmid DNA (pDNA). Log2-transformed normalized read counts were scored for gene effect by averaging the fold change for constructs sharing the same target relative to either pDNA or DMSO read counts. A z-score was then assigned to each gRNA construct and to each target gene, the latter based on average log (fold change) relative to pDNA or DMSO samples. gRNA constructs with low abundance (z ≤ -3) in the DMSO condition were excluded from analysis. Statistical significance per target was calculated from target gene z-scores using the standard distribution of the average log(fold changes). Average log(fold change) and -log10(p-value) were plotted for genes with three or greater constructs remaining.

### GeoMx-based spatial transcriptomics

Spatial transcriptomic profiling of the NC TMA was performed on the GeoMx Digital Spatial Profiler (DSP) (NanoString) as described^75^ at the Dana-Farber Cancer institute (DFCI) Pathology Core. Immunofluorescence was performed to identify macrophages (anti-CD68, clone KP1, catalog no. sc- 20060, Santa Cruz) and all immune cells (anti-CD45, clone 2B11+PD7/26, catalog no. NBP2-34528, Novus Biologicals, Centennial, CO). Nuclei were stained using SYTO13 (Thermo Fisher Scientific). Tumor cells were identified amongst non-immune populations using large nuclear density and size-threshold settings in consultation with a pathologist for all cores (C. French). Based on the stained cells and tissue morphology, regions of interest (ROI, ≥ 50 cells) were digitally selected on slides that have been pre-hybridized with barcoded-oligonucleotide probes complementary to target RNA using the Cancer Transcriptome Atlas (CTA) oligo library (1,822 genes). The DNA barcodes within the ROI were cleaved off by UV light and sent for RNAseq at the Molecular Biology Core Facilities (MBCF) at DFCI.

Following the collection of cleaved barcodes on the GeoMx DSP instrument, the 96-well collection plate was submitted to the Molecular Biology Core Facilities at Dana-Farber Cancer Institute where the collection plate was dried down using a permeable membrane (Sigma, A9924) and resuspended in 10 µl of molecular biology grade nuclease-free water. PCR amplification was performed using 4 µl per ROI according to the GeoMx DSP NGS Readout User Manual. Uniquely dual-indexed libraries were pooled and quantified by Qubit fluorometer and Agilent TapeStation 2200. Sequencing was performed on the NovaSeq 6000 instrument using an S1 flow cell with paired-end 27 bp reads. Demultiplexing and FASTQ file generation were performed using Illumina bcl2fastq v2.20 software. FASTQ files were processed further using the geomxngspipeline v2.0.0.16 software to generate DCC files.

### Analysis of GeoMx spatial transcriptomics

All tumor-compartment segments from the five TMA slides were checked with default quality control parameters using the GeoMx DSP Analysis Suite (n = 203). Segments with a nuclei count < 400 were dropped from further analysis to eliminate segments with nuclei counts outside of the manufacturer’s recommended ten-fold range and avoid comparisons between segment counts with extreme differences in size.

Low outlier probes were removed for which the average counts across all segments were ≤ 10% of the counts for all probes for that gene. Additionally, all probes that failed the Grubbs outlier test in ≥ 20% of segments were removed. Using the default GeoMx Analysis parameters, biological probes were collapsed to gene-level counts.

Gene-level counts from the remaining 173 segments were normalized to the geometric mean of 32 housekeeping genes preselected in the GeoMx analysis suite. Normalized mRNA count distribution quartile ranges were then visualized per segment and 19 segments with poor signal differentiation were removed. Because of our interest in the expression of CDKN2A and EZH2, the segments were removed where the signal-to-noise ratio of either CDKN2A or EZH2 mRNA was below detection. The detection level was defined as a 2-fold greater expression compared with CTA negative probe counts. Correlation coefficients between six genes of interest were calculated from the log counts of the six genes using the selected23 segments which expressed both CDKN2A and EZH2.

### siRNA knockdown

The following siRNAs were purchased from Dharmacon (Lafayette, CO): siCTRL, ON-TARGET plus siRNA #1 (Dharmacon; cat no. D-001810-01-20); si*NUT* (targeting coding sequence), AAACUCAGAACUUUAUCCUUAUU. siRNA was transfected using Lipofectamine RNAiMAX Transfection Reagent (13778150, Thermo Fisher Scientific) at a final concentration of 50 nM per siRNA.

### Cell Cycle Analysis by Flow Cytometry

2x10^^6^ cells were trypsinized and spun at 500 x g for 5 minutes at 4 °C, and washed twice with PBS. Pellets were resuspended in 1 ml of cold PBS and added dropwise while gently vortexing to 9 ml 70 % ethanol in a 15 ml polypropylene centrifuge tube. Fixed cells were then frozen at –20 °C overnight. The next day, cells were centrifuged at 500 x g for 10 minutes at 4 °C and washed twice with 3 ml of cold PBS. Cells were resuspended in 400 µl of propidium iodide staining solution (0.2 mg/ml RNAse A, 0.02 mg/ml propidium iodide, 0.1 % Triton-X in PBS) and incubated for 15 minutes at 37 °C. Samples were then transferred to ice and analyzed on a BD LSR II flow cytometer at the Harvard Medical School flow cytometry core (https://immunology.hms.harvard.edu/resources/flow-cytometry). Cell cycle analysis was performed using FCS Express 7 (De Novo Software, Pasadena, CA).

### Procedures and analysis to evaluate in vitro synergy of compound combinations

To investigate synergy with the combination of two compounds, the concentration of one is fixed at 0 or 25% of its IC50, while the other is varied to re-measure its IC50s. The reciprocal procedure is performed. The data enables a calculation of the combination indices using Chou-Talalay Drug combination studies and their synergy quantification using the Chou-Talalay methods with free online software (Compusyn) to determine synergy, additivity, or antagonism.

### Cleavage Under Targets & Release Using Nuclease (CUT&RUN)

NC cells and *Drosophila* Kc167 spike-in cells were harvested at room temperature, resuspended in Wash Buffer (20 mM HEPES-NaOH pH 7.5, 150 mM NaCl, 0.5 mM spermidine, 1× Roche protease inhibitor cocktail), and counted using a Vi-CELL Blu cell counter. For each antibody, 400,000 NC cells were mixed with 100,000 Kc167 cells then incubated for 5-10 minutes with 10 µl of Concanavalin A-coated magnetic beads (Bangs Laboratories #BP531) in Binding Buffer (20 mM HEPES-KOH pH 7.9, 10 mM KCl, 1 mM CaCl2, 1 mM MnCl2). Bead-bound cells were permeabilized in 50 µL Antibody Buffer (20 mM HEPES-NaOH pH 7.5, 150 mM NaCl, 0.5 mM spermidine, 0.01% digitonin for 10-15 cells or 0.0025% digitonin for PER-403 cells, 2 mM EDTA, 0.1% BSA, 100 nM TSA, 0.1 unit/ml citrate synthase, 1 mM Oxaloacetic acid, 1× Roche protease inhibitor cocktail). For negative control samples (e.g. rabbit α-mouse IgG) as well as H3K27me3 samples 1 µL of SNAP-CUTANA K-MetStat Panel (Epicypher #19-1002) was added to each sample. Samples were then incubated with 1 µL of anti-H3K27ac antibody (Cell Signaling Technologies #8173), anti-H3K27me3 antibody (Cell Signaling Technologies #9733), or rabbit α-mouse IgG (abcam #ab46540) overnight at 4°C with nutation. Bead-bound cells were washed twice with 200 µL Wash Buffer + 0.01% digitonin for 10-15 cells or 0.0025% digitonin for PER-403 cells, and then incubated with 700 ng/ml pA-MNase in 100 µL Wash Buffer + 0.01% digitonin for 10-15 cells or 0.0025% digitonin for PER-403 cells for 1 h at 4°C with nutation. Bead-bound cells were washed twice in 200 µL Wash Buffer + 0.01% digitonin for 10-15 cells or 0.0025% digitonin for PER-403 cells. For 10-15 cells, cells were resuspended in 200 µL low-salt Wash Buffer (20 mM HEPES-NaOH pH 7.5, 0.5 mM spermidine, 0.01% digitonin, 1x Roche protease inhibitor cocktail), the pA-MNase activated for 15 minutes at 0°C with 100 µL ice-cold Calcium Incubation Buffer (3.5 mM HEPES-NaOH pH 7.5, 10 mM CaCl2, 0.01% digitonin), the reaction stopped and chromatin fragments were released by exchanging the beads into 100 µL Stop Buffer (170 mM NaCl, 20 mM EGTA, 0.01% digitonin, 50 µg/ml RNase A) followed by incubation for 30 min at 37°C. For PER-403 cells, cells were resuspended in 50 µL Wash Buffer + 0.0025% digitonin, 1 µL 100 mM ice-cold CaCl2 was added to activate the pA-MNase followed by incubationg the cells for 30 minutes at 0°C, the reaction stopped by adding 50 µL of STOP buffer (340 mM NaCl, 20 mM EDTA, 4 mM EGTA, 0.0025% digitonin, 50 µg/mL glycogen, 100 µg/mL RNase A), and chromatin fragments were released by incubating for 30 min at 37°C. To extract the DNA, regardless of cell type, 200 µl of Oligo Binding Buffer (Zymo) followed by 800 µL of 100% ethanol was added to each sample and the total volume was then loaded onto a Zymo-Spin DCC-5 by centrifugation. The column was washed twice with 200 µL Zymo DNA Wash Buffer and then the DNA was eluted from the column in 15 µl of DNA Elution Buffer. Sample concentration was measured using a DeNovix High Sensitivity dsDNA Assay on a DeNovix DS-11 FX+ Fluorometer. Illumina libraries were prepared using a NEBNext Ultra II Library Prep Kit for Illumina with NEBNext Multiplex Oligos for Illumina with the following modifications: All volumes were reduced 2-fold. The NEBNext Adaptor was diluted 1:25 in NEBNext Adaptor Dilution Buffer. After Adaptor Ligation, a DNA cleanup using 1.0x volumes of Sera-Mag Select beads (Cytiva #29343052) was conducted. The PCR thermal cycling conditions were as follows: 1 cycle: 98°C for 45 s, 15 cycles: 98°C for 15 s, 65°C for 10 s, 1 cycle: 72°C for 1 min, hold: 4°C. After PCR amplification, DNA cleanup was conducted twice with 1.0x volumes of Sera-Mag Select beads. Quality control was conducted on the resulting 15 μl eluate using an Agilent 4200 TapeStation D1000 ScreenTape to determine sample concentration and sample quality. Libraries were sequenced on an Illumina NextSeq 2000 to generate 50 bp paired-end reads.

### Confocal microscopy procedures and analysis

10-15 and PER-403 cells were seeded at a density of 300,000 cells per well on #1.5 cover glasses (VWR, 16004-326) in 6-well format. Samples were fixed with 4% paraformaldehyde for 5 minutes and blocked in 5% goat normal serum (Cell Signaling Technologies, 5425) for an hour before antibody detection using antibodies listed in Supplementary Tables S5-S6, according to manufacturer’s protocol. Samples were mounted onto coverslips (Fisherbrand, 12-550-15) with SlowFade glass antifade mountant (Invitrogen, S36917). Images were acquired on a Zeiss Axio Observer Z1 single point scanning confocal microscope with a linear encoded motorized stage from Zeiss, a LSM 980 scan head and Toptica iChrome MLE laser CW launch using the Airyscan2 super-resolution external detector. A Zeiss Plan Apo 63x/1.4 DIC III oil immersion objective lens was used for the imaging. Images were acquired using a galvo, unidirectional scan with a pixel dwell time of 0.75us, 2x line averaging, and a pixel size of 42nm x 42nm (set as optimal for Airyscan detection) by Zen Blue Acquisition Software and exported as CZI files. For all images pinhole diameter was 5AU (set as optimal for Airyscan detection). Alexa Fluors were excited with either 493nm or 590nm solid state diode lasers with AOTF modulation, and emissions were collected on a GaAsP-PMT detector with collection windows of 380-608nm and 527-735nm, respectively. Super Resolution images were generated by Airyscan SuperResolution 2D automatic processing (Weiner noise filtering value range 6.4-7.5 as optimally determined by the Zen software). Laser power and detector settings for each sample are listed in Supplementary Tables S7- S8. Files were converted using Arivis SIS Converter (ZEISS). Pearson correlation of fluorescent signals was performed on twenty-four nuclei per condition using Arivis PRO Image Analysis Software (Zeiss). Images cropped to isolate individual nuclei and the Pearson value for all voxels was recorded. Resulting values were analyzed in GraphPad Prism. 2D fluorescence intensity profiles for each channel were generated using Fiji ImageJ.

### Xenograft efficacy and pharmacodynamic studies

All *in vivo* studies were conducted at Dana-Farber Cancer Institute with the approval of the Institutional Animal Care and Use Committee in an AAALAC accredited vivarium. NMC models were established by injecting 1 x 10^6^ PER403, 10-15 cells or 2 x 10^6^ 14169 cells in 7-week old female NSG mice obtained from Jackson Laboratory (Bar Harbor, ME). Bioluminescent imaging was performed to verify disseminated tumor establishment in mice by injecting D-luciferin subcutaneously at 75 mg/kg (Promega) and imaged with the IVIS Spectrum Imaging System (Perkin Elmer). To quantify bioluminescence, identical regions of interest were drawn and the integrated total flux of photons (the sum of the prone and supine values) using the Living Image software (Perkin Elmer) were used for initial randomization of mice into various treatment groups and subsequently weekly imaging for assessing tumor response. Mice implanted with PER403 and 10-15 cells were randomized 7 days after cell implantation, while mice implanted with 14169 cells were randomized 20 days after cell implantation.

ABBV-075 and ABBV-744 compounds were formulated in 2% DMSO, 30% PEG 400, and 68% Phosal-50PG, and administered once daily by oral gavage for 28 days. Taz was formulated in 0.5% Methylcellulose (400cP) + 0.1% Tween 80 in water, pH4.0, and administered by oral gavage twice daily for 28 days. Vehicle-only treatments (2% DMSO, 30% PEG 400, and 68% Phosal-50PG) were administered once daily by oral gavage. Bioluminescent imaging was performed once weekly after treatment initiation and body weights were measured twice weekly.

For pharmacodynamic (PD) studies, tumor bearing NSG mice were treated with either vehicle control (2% DMSO, 30% PEG 400, 68% Phosal-50PG), ABBV-744 (37.5 mg/kg, once daily) or taz (250 mg/kg, twice daily) administered for 5 days. After the last dose, three animals per group were euthanized at 2 hours and tumor samples (e.g. ovaries, liver, brain) were collected and fixed in 10% buffered formalin for immunohistochemistry.

To compare survival statistically, log-rank (Mantel-Cox) test was used. To compare MYC and Ki-67 counts, unpaired t-test was used.

## Accession numbers

RNA-seq data will be available in Gene Expression Omnibus (GEO) with accession number GSE142481. CUT&RUN data will be available in GEO with accession number GSE228533.

## ACKNOWLEDGEMENTS

ABBV-075 and ABBV-744 were generously provided by AbbVie, Inc.. PZ-6438 (tazemetostat) and EPZ- 5676 (pinometostat) were provided by Epizyme (an Ipsen company). Ipsen reviewed this manuscript for scientific accuracy, but had no input into the content.

