## Supplementary Figures for "EZH2 synergizes with BRD4-NUT to drive NUT carcinoma growth through silencing of key tumor suppressor genes"

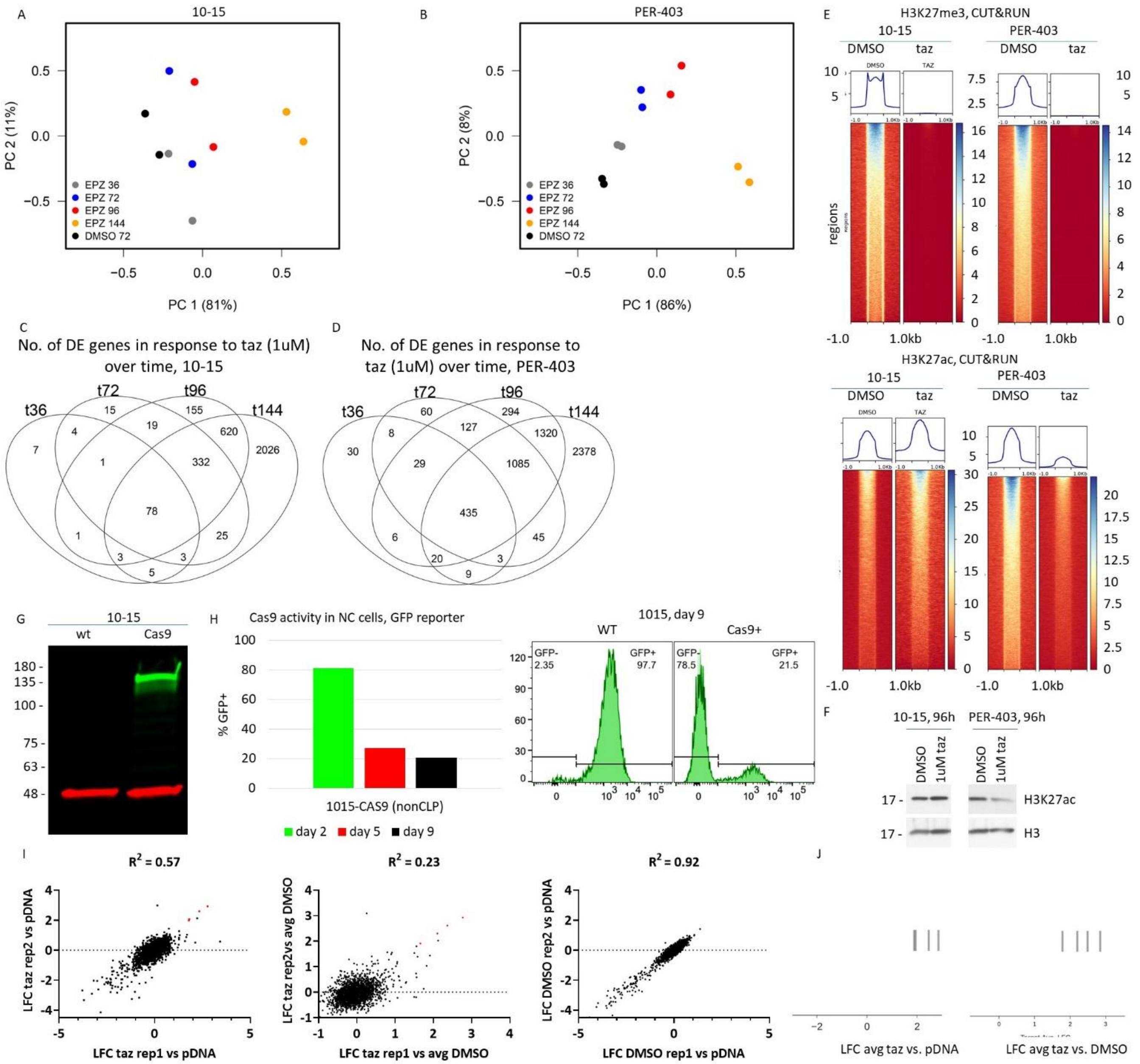


**Supplementary Fig. S1.** Genes repressed by EHZ2-induced H3K27 trimethylation include *CDKN2A*, a key NC tumor suppressor. A-B. Principal component analysis of RNAseq based on the top 500 genes by variance for the four different time points indicated. EPZ 36, EPZ-6438 (taz, 1µM) treatment for 36h; EPZ 72, 96, 144, same for 72h, 96h, and 144h; DMSO 72, DMSO treatment for 72h. C-D. Petal plots comparing DE genes identified by RNAseq at the time points indicated. E. Heat maps of CUT&RUN in cells treated with taz for 96h or DMSO 72h corresponding with the experiment in A-D. F. Immunoblot of extracted histones as indicated. G. Immunoblot comparing wild type with the CAS9-transduced derivative. H. Flow cytometric analysis of GFP-positive fractions over time. I. Replicate reproducibility of gene effect using DMSO-treated or pDNA as baseline. Red colored dots represent the four gRNAs targeting *CDKN2A*. J. Rug plot demonstrating individual effects of each *CDKN2A* gRNA.

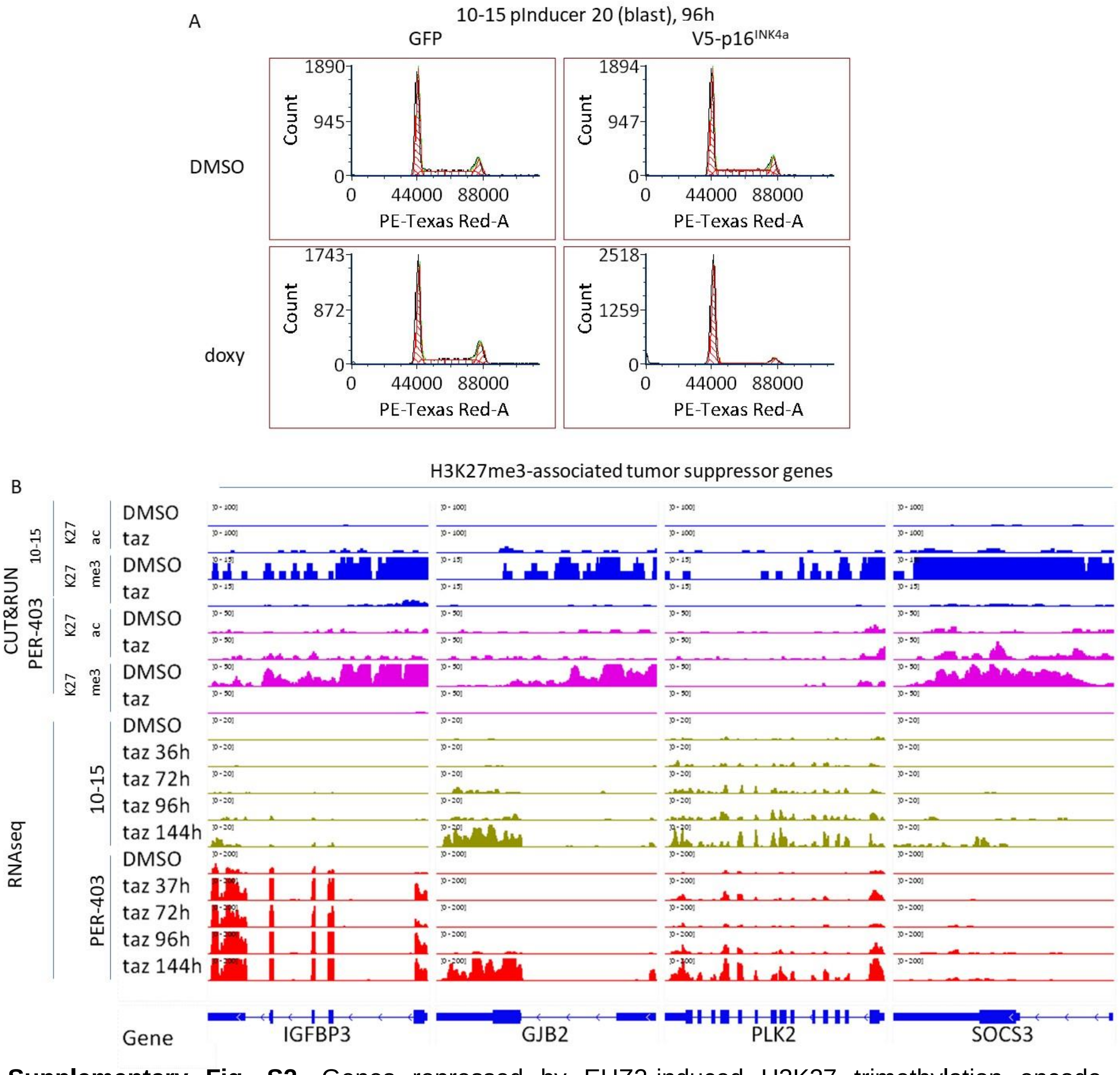


**Supplementary Fig. S2.** Genes repressed by EHZ2-induced H3K27 trimethylation encode numerous tumor suppressors, including *CDKN2A*. A. Flow cytometric cell cycle analysis tracings of single representative replicate, of two biologic replicates, corresponding with experiment in 3C. B. Integrated genome viewer views CUT&RUN and RNAseq peaks at the indicated genes. Each track shown is from one of two biologic replicates. K27ac, CUT&RUN using anti-histone H3 K27ac; K27me3, CUT&RUN using anti-histone H3 K27me3.

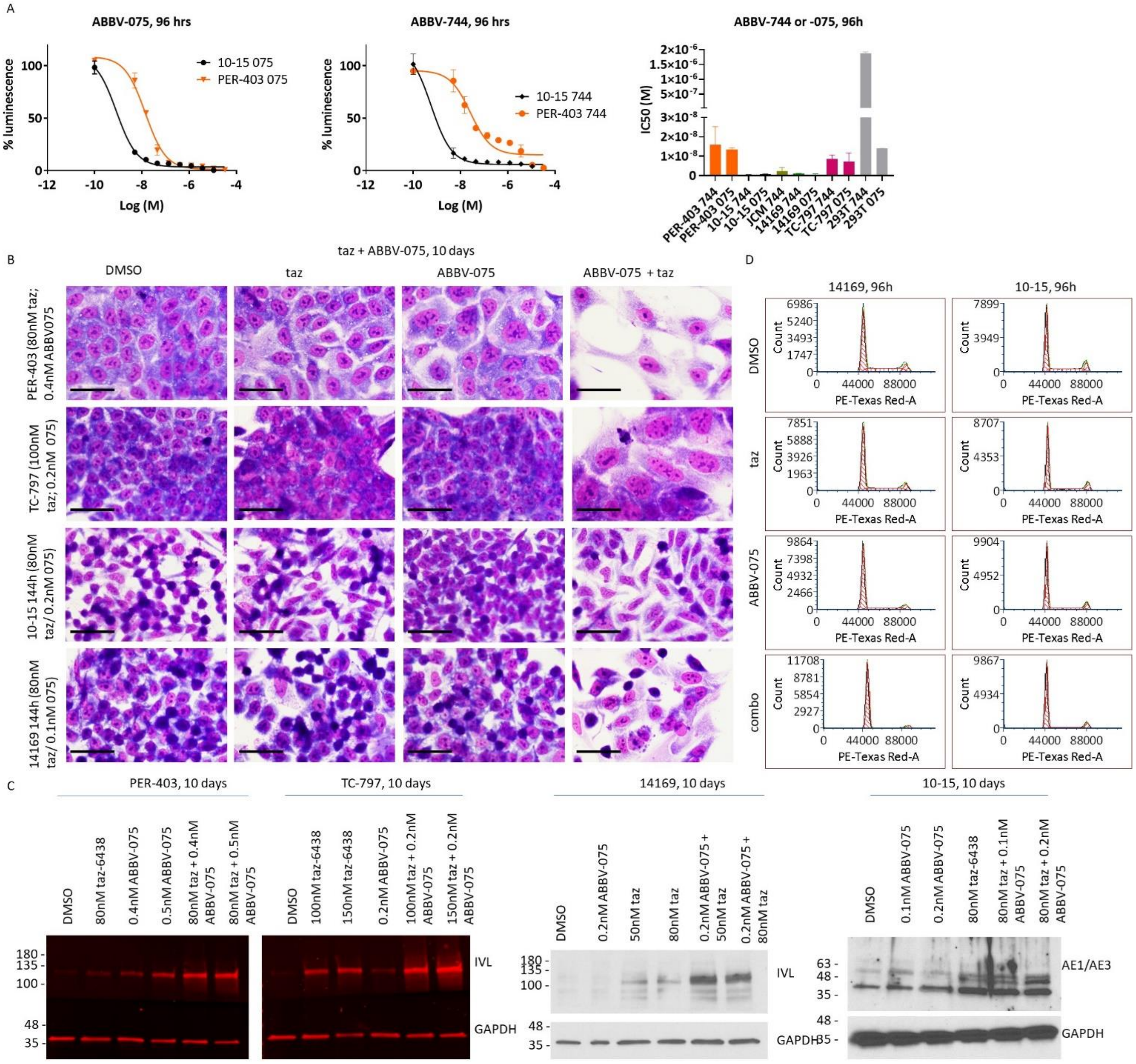


**Supplementary Fig. S3.** Combined EZH2 and BET inhibition synergistically induces terminal differentiation and blocks growth of NC. A. Left, dose response curves from representative single replicates using Cell Titer Glo as readout. Right, IC50s from biological triplicate, technical triplicate, dose-response assays corresponding with the experiments depicted on the left. 744, ABBV-744; 075, ABBV-075. B. Hemacolor stained cells grown on coverslips. Scale bar, 25µm. C. Immunoblots as indicated. D. Flow cytometric cell cycle analysis tracings of single representative replicate, of two biologic replicates, corresponding with experiment in 5E.

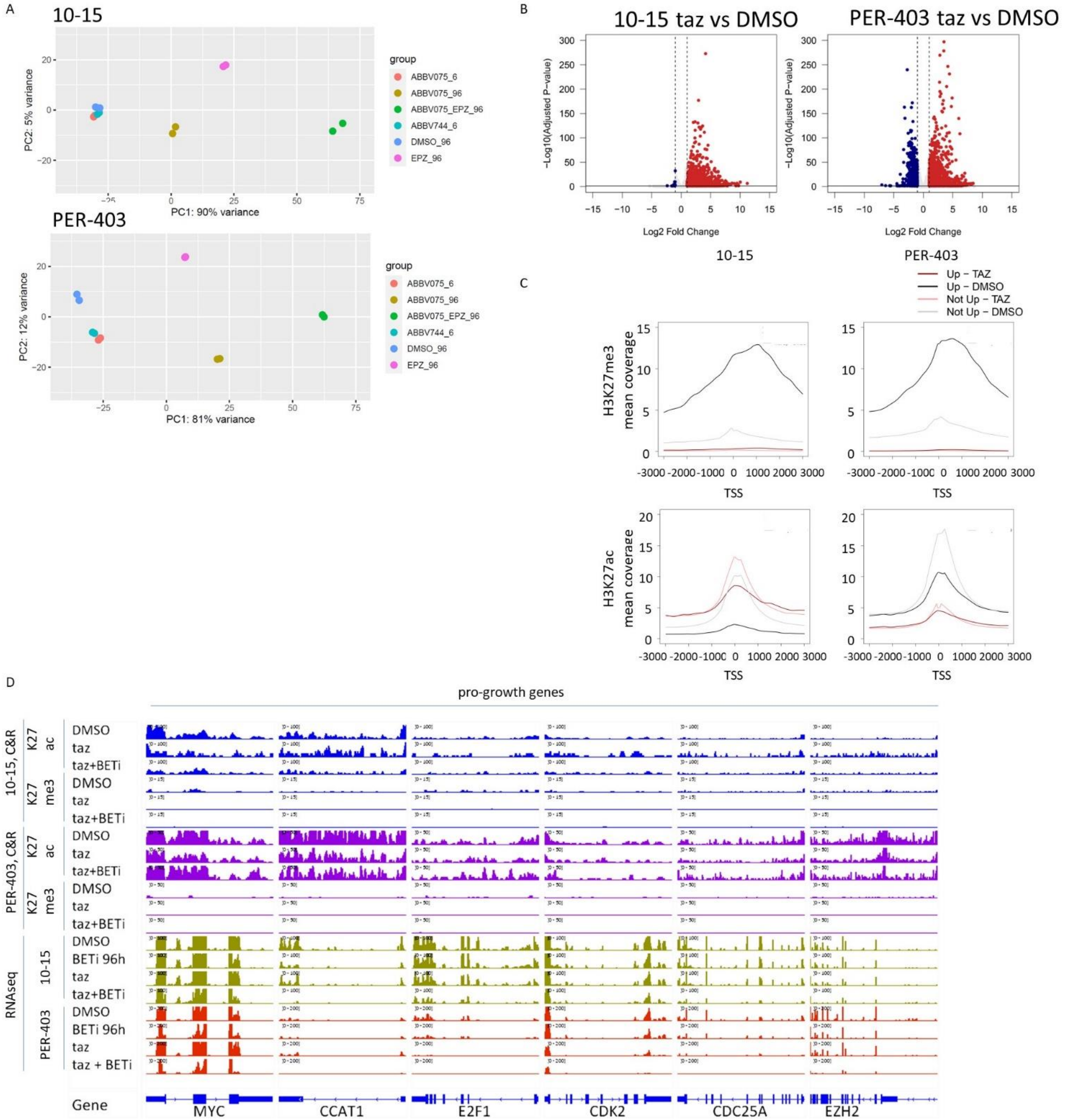


**Supplementary Fig. S4.** Combined BET and EZH2 inhibition synergizes to downregulate cell proliferation genes. A. Principal component analysis of RNAseq on cells treated as indicated for 6h or 96h, based on top 500 genes by variance. B. Volcano plots of DE genes identified by RNAseq comparing DMSO-treated with taz-treated cells for 96h. Data corresponds with that in A. C. Enrichment profiles in H3K27me3- and H3K27ac-associated chromatin at the transcriptional start site (TSS) of coding genes. D. Integrated genome viewer views CUT&RUN and RNAseq peaks at the indicated genes. Each track shown is from one of two biologic replicates.

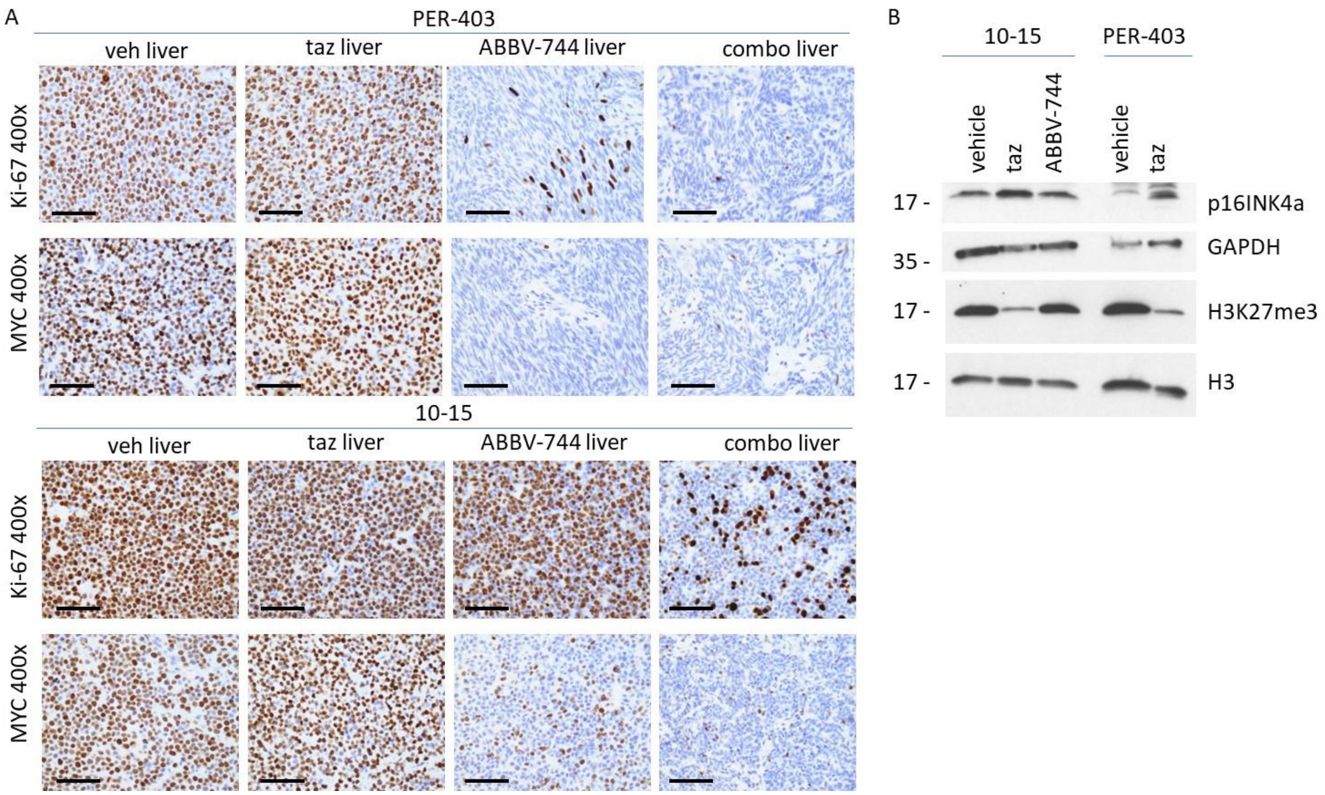


**Supplementary Fig. S5.** Pharmacodynamic analysis demonstrates cooperative effects of combined EZH2i and BETi on cell proliferation. A. Representative images of MIB-1 (Ki-67) and MYC staining used to quantify percent cells staining depicted in Fig. 7E. Scale bars, 50µm. B. Immunoblots from single, frozen, harvested metastatic tumors to ovary, each from an individual mouse. Only tumors replacing the entire ovary were used to avoid contaminating normal ovarian tissue.
