## Supplementary Table S2-S3 for "EZH2 synergizes with BRD4-NUT to drive NUT carcinoma growth through silencing of key tumor suppressor genes"

| Supplementary Table S2. antibodies used for immunohistochemistry. | | | | | | |
|---|---|---|---|---|---|---|
| protein recognized | dilution | catalogue no. | clone | company | detection | antigen retrieval |
| cleaved PARP | 1:50 | 5625 | (ASP214) D64E1 | CST* | Leica Bond – Leica Refine Detection Kit (DS9800) | citrate |
| Ki-67 | 1:500 | M7240 | MIB-1 | Dako | Leica Bond – Leica Refine Detection Kit (DS9800) | EDTA |
| MYC | 1:100 | ab32072 | Y69 | Abcam | Leica Bond – Leica Refine Detection Kit (DS9800) | EDTA |
| p16INK4a | 1: 60,000 | ab54210 | 2D9A12 | Abcam | Leica Bond – Leica Refine Detection Kit (DS9800) | Citrate |
| keratin cocktail | 1:200 | M3515 | AE1/AE3 | Dako | Leica Bond – Leica Refine Detection Kit (DS9800) | Citrate |
| H3K27me3 | 1:100 | 9733 | C36B11 | CST | Leica Bond – Leica Refine Detection Kit (DS9800) | Citrate |
| EZH2 | 1:100 | 5246 | D2C9 | CST | Leica Bond – Leica Refine Detection Kit (DS9800) | EDTA |
| NUT | 1:100 | 3625 | C52B1 | CST | Leica Bond – Leica Refine Detection Kit (DS9800) | |

*Cell Signaling Technologies

| Supplementary Table S3. antibodies used for immunoblotting. | | | | |
|---|---|---|---|---|
| protein recognized | dilution | catalogue no. | clone | company |
| CAS9 | 1:1,000 | 632607 | TG8C1 | Takara Bio |
| AE1/AE3 | 1:500 | 313M | AE1/AE3 | Millipore-Sigma |
| KRT7 | 1:1,000 | 4465 | D1E4 | CST |
| H3K27me3 | 1:1,000 | 9733 | C36B11 | CST |
| V5 tag | 1:1,000 | 13202 | D3H8Q | CST |
| BIRC5 | 1:1,000 | 2808 | 71G4B7 | CST |
| Phospho-Rb (Ser780) | 1:1,000 | 9307 | rabbit polyclonal | CST |
| RB | 1:500 | sc-102 | IF8 | Santa Cruz Bio |
| p16INK4a | 1:1,000 | 10883-1-AP | rabbit polyclonal | Proteintech |
| GAPDH | 1:20,000 | AM4300 | 6C5 | Invitrogen |
| involucrin | 1:1000 | I9018 | SY5 | Millipore Sigma |
| H3K27ac | 1:2,500 | 4353 | rabbit polyclonal | CST |
| histone H3 | 1:20,000 | ab1791 | rabbit polyclonal | Abcam |
| NUT | 1:1,000 | 3625 | C52B1 | CST |
| PARP | 1:1,000 | 9542 | rabbit polyclonal | CST |
