## Supplementary Tables S5-S8 for "EZH2 synergizes with BRD4-NUT to drive NUT carcinoma growth through silencing of key tumor suppressor genes"

| Supplementary Table S5. Primary antibodies used for immunofluorescence confocal microscopy. | | | | |
| --- | --- | --- | --- | --- |
| protein recognized | dilution | catalogue no. | clone | company |
| H3K27me3 | 1:1,000 | 9733 | C36B11 (rabbit) | CST |
| H3K27ac | 1:2,500 | 4353 | rabbit polyclonal | CST |
| H3K27ac | 1:1,000 | 39685 | MABI 0309 (mouse) | Active Motif |
| NUP62 | 1:400 | 24588 | E4Y5P (rabbit) | CST |

| Supplementary Table S6. Secondary antibodies used for immunofluorescence confocal microscopy. | | | |
| --- | --- | --- | --- |
| Species recognized/Wavelength | dilution | catalogue no. | company |
| Anti-mouse Alexa Fluor 488 | 1:2,000 | **A-11001** | Invitrogen |
| Anti-mouse Alexa Fluor 594 | 1:1,000 | A-11005 | Invitrogen |
| Anti-rabbit Alexa Fluor 488 | 1:2,000 | A-11008 | Invitrogen |
| Anti-rabbit Alexa Fluor 594 | 1:1,000 | A-11012 | Invitrogen |

| Supplementary Table S7. 10-15 laser power and detector settings | | | | | | |
| --- | --- | --- | --- | --- | --- | --- |
|  | Test | | Positive Control | | Negative Control | |
| Channel | 488 | 594 | 488 | 594 | 488 | 594 |
| Laser Power (%) | 1.5 | 0.9 | 0.1 | 0.1 | 0.1 | 4.0 |
| Digital Gain (V) | 700 | 625 | 725 | 700 | 650 | 700 |
| Gain | 2 | 2 | 3 | 3 | 4 | 5 |
| Offset | 0 | 0 | 0 | 0 | 0 | 0 |

| Supplementary Table S8. PER-403 laser power and detector settings | | | | | | |
| --- | --- | --- | --- | --- | --- | --- |
|  | Test | | Positive Control | | Negative Control | |
| Channel | 488 | 594 | 488 | 594 | 488 | 594 |
| Laser Power (%) | 0.8 | 0.08 | 0.3 | 0.05 | 0.09 | 1.5 |
| Digital Gain (V) | 650 | 660 | 650 | 700 | 700 | 740 |
| Gain | 5.5 | 5.2 | 5.0 | 5.5 | 4.5 | 4.2 |
| Offset | 0 | 0 | 0 | 0 | 0 | 0 |
